# Chimeric Immunoglobulin and human Immunoglobulin M structures provide insights on joining-chain independent assembly and function

**DOI:** 10.1101/2025.07.09.663956

**Authors:** Mengfan Lyu, Beth M. Stadtmueller

**Affiliations:** Department of Biochemistry, University of Illinois Urbana-Champaign, Urbana, Illinois 61801 USA; Department of Biomedical and Translational Sciences, Carle Illinois College of Medicine, University of Illinois Urbana-Champaign, Urbana, Illinois 61801 USA; Carl R. Woese Institute of Genomic Biology, University of Illinois Urbana-Champaign, Urbana, Illinois 61801 USA

## Abstract

Polymeric (p) immunoglobulins (Igs) play critical roles in vertebrate immunity. IgM is the evolutionarily oldest pIg and functions both in circulation and in the mucosa. pIgM typically comprises between four and six IgM monomers and up to one joining chain (JC), which in mammals facilitates pIg assembly and promotes delivery to mucosal secretions. Bony fish (teleosts) lack JC and assemble tetrameric IgM whereas humans can express JC-containing pentamers and JC-free hexamers. Here we report cryo-electron microscopy structures of two JC-free chimeric IgM, comprising bony fish and human sequences, and the structure of human hexameric IgM. Chimeric IgM structures adopted unique pentameric geometry distinct from both human and fish pIgM whereas the human hexameric IgM structure adopted hexagonal geometry similar to JC-containing pentameric IgM, albeit with structural differences in center of the molecule. Together results provide new insights on how IgM heavy chain motifs contribute to JC-free pIgM assembly and reveal plasticity of this process, which can be manipulated to create pIg structures not observed in nature. Moreover, we found that antigen-targeting chimeric IgM could neutralize *C. difficile* toxin cytotoxicity, indicating potential to engineer uniquely structured pIgs to prevent or treat disease.

## INTRODUCTION

Immunoglobulin (Ig) M is a primordial Ig subclass present in essentially all jawed vertebrates (gnathostomes). It is the first antibody produced during the humoral immune response and serves important functions in complement system activation and adaptive immunity^1,2^. In most species IgM adopts polymeric (p) structures; only cartilaginous fish are known to express functional, monomeric (m) IgM^3,4^. Whereas the domain organization and structural folds of mIgM are evolutionarily conserved (see Fig. 1a), structural organization of pIgM varies among and within species. Human IgM pentamer (hereafter IgM-J) consists of five covalently linked mIgM and one 15-kDa protein named joining chain (JC); alternatively, human IgM forms JC-free hexamers which exist at low abundance and make up 5-10% of the circulatory IgM^5–7^. In lower jawed vertebrates (cartilaginous fish, teleosts, amphibians, and reptiles), pIgM consists of four to six IgM monomers and up to one JC, except for teleosts, which have lost the JC during evolution and produce JC-free IgM tetramers^4^. IgM is the first antibody expressed by B cells during the immune response and generally has a lower affinity for antigen due to lack of affinity maturation but affords higher avidity as compared to monomeric Ig subclasses. Moreover, the multivalent nature makes pIgM more efficient at complement activation and inducing complement dependent cytotoxicity; a recent study found that JC-deficient hIgM hexamer outcompetes both hIgM pentamer and IgG monomer at low antigen density and/or concentration^8,9^.

**Fig. 1.**
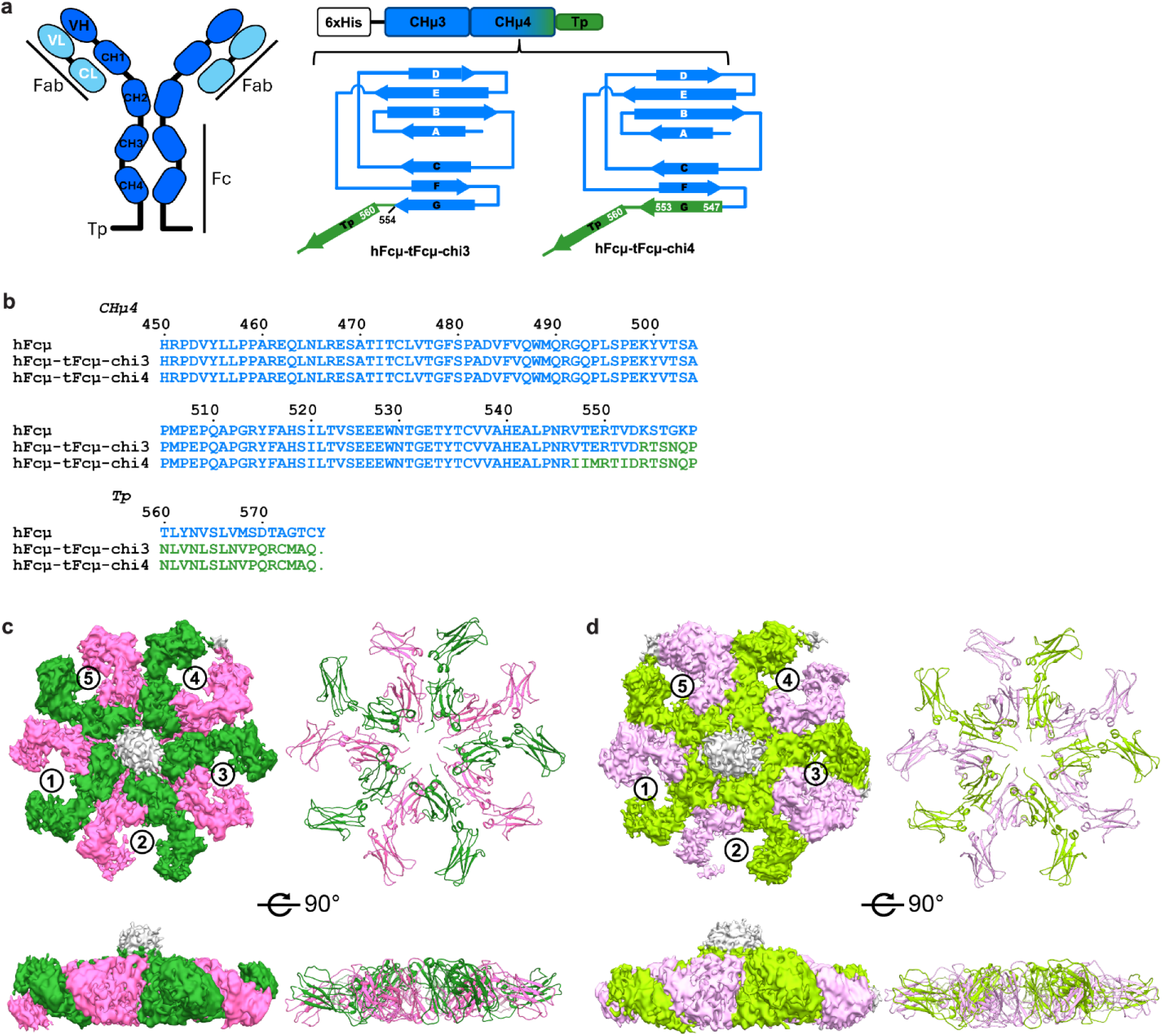
Construct design and structural overview of hFcμ-tFcμ-chi3 and hFcμ-tFcμ-chi4. **a** Schematic diagrams of an IgM monomer (*left*) and the cFcμ construct design (*right*). The IgM monomer is labeled to indicate: VHμ, variable domain of heavy chain, VL, variable domain of light chain, CHμ, constant domain of heavy chain, and CL, constant domain of light chain. The schematic showing cFcμ design includes domain organization on the top and CHμ4 domain topology below. Each cFcμ sequence consists of an N-terminal hexa-histidine tag, partial hFcµ sequence (blue), and tFcµ C-terminal sequence (green). **b** Sequence alignment between hIgM heavy chain, hFcμ-tFcμ-chi3, and hFcμ-tFcμ-chi4. The hIgM heavy chain sequence is colored blue and the tIgM sequence is colored green. **c** hFcμ-tFcμ-chi3 cryo-EM density map (*left*) and structure (cartoon representation, *right*) shown in two orientations. **d** hFcμ-tFcμ-chi4 cryo-EM density map (*left*) and structure (cartoon representation, *right*) shown in two orientations. In **c** and **d**, individual chains are colored pink (chains A, C, E, G, K) or green (chains B, D, F, H, L) and the Fcµ monomers are numbered. Map regions corresponding to unmodeled density are shown in gray.

In addition to being released into circulation, pIgM can also be transported to the mucosa by binding to the polymeric Ig receptor (pIgR), which is expressed on the basolateral surface of epithelial cell barriers. The pIgM is transcytosed to the apical surface of epithelial cells where pIgR undergoes proteolytic cleavage, releasing its ectodomain, named secretory component (SC), bound to pIgM. The complex consisting of SC and pIgM is secreted into the mucosa and referred to as secretory (S) IgM^10^. The pIgR-mediated IgM secretion mechanism is conserved in jawed vertebrates despite structural divergence in the SC and pIgM structure across species^10,11^. SIgM, though less abundant than SIgA in humans, provides first-line defense in mucosal secretions through mechanisms including antigen agglutination and crosslinking, and may play a larger role in lower gnathostomes lacking a specialized secretory Ig such as SIgA in mammals^12,13^. Human SIgM-antigen complexes can also be reverse transcytosed across the mucosal cell barriers by FcμR/TOSO, suggesting a role in antigen presentation and/or other immune functions^14^.

SIgM and pIgM domain organization is similar among species; however structural variations have been reported. IgM monomer structure is highly conserved, each containing two heavy chains comprising one variable domain VHμ and four constant domains, CHμ1-CHμ4, as well as two light chains comprising one variable domain VL1 and one constant domain CL1. Together these domains form two antigen binding fragments (Fabs) and one constant, or crystallizable, fragment (Fcμ; see Fig. 1a). A 17-residue C-terminal extension called tailpiece (Tp) follows CHμ4. Prior cryogenic electron microscopy (cryo-EM) studies resolved multiple structures containing a pentameric IgM-J Fcμ core (Fcμ-J), pIgM lacking CHμ2 and Fabs. The human SFcμ-J (Fcμ-J bound to SC) structures revealed an asymmetric arrangement of five Fcμ monomers (Fcμ1-5) linked through ten Tps, with JC occupying the gap between Fcμ1 and Fcμ5 and SC bound to one face of the Fcμ-J^15,16^. At the center of Fcμ-J, C-terminal Tps formed two anti-parallel β-sheets joined by JC β-strands on both faces; JC also mediated binding between SC and hIgM-J pentamer. A subsequent structure of full-length human IgM-J pentamer revealed a flexible region between the second and third heavy chain constant domains (CHμ2 and CHμ3) that allows Fab and CHμ2 domains to pivot together as a subunit^17^. Cryo-EM structures of hexameric IgM remain unreported; however a hexagonal model assembled from multiple data types has been published^18,19^.

In 2023, we reported a 2.8Å resolution structure of the teleost (t) IgM Fc (tFcμ), containing four Fcμ monomers (Fcμ1-Fcμ4)^20^. In the JC-free structure, four Fcμ monomers were linked through eight Tps which form a β-sandwich assembly situated between Fcμ1 and Fcμ4. Notably, two of the eight heavy chains in tFcμ adopted distinctive folding patterns, giving rise to a structural configuration that markedly diverged from structures containing Fcμ-J^15^. We also identified C-terminal sequences associated with structural features unique to tFcμ and showed that when combined with human IgM CHμ3-CHμ4 sequences, teleost sequences could promote JC-free, polymeric assembly of chimeric (c) Fcμ^20^. Here we report the cryo-EM structures of two cFcμs and JC-free hIgM. These three structures share similarities and differences with published Fcμ-J containing structures and the tFcμ structure relevant to understanding JC-free pIgM assembly in multiple species. Furthermore, we report that cFcμs can be engineered to contain single domain antibodies thereby providing a structurally unique scaffold to target disease antigens.

## RESULTS

### Cryo-EM structures of hFcμ-tFcμ-chi3 and hFcμ-tFcμ-chi4 reveal JC-free pentamers

Previously, we reported the structure of tFcμ and structure-based design of cFcμ, in which we replaced 22-37 C-terminal residues of human IgM heavy chain with homologous counterparts from the tIgM heavy chain sequence^20^. Here we determined the cryo-EM structures of two cFcμs, hFcμ-tFcμ-chi3 and hFcμ-tFcμ-chi4, which include tIgM residues 426-448 and tIgM residues 420-448, respectively (Fig. 1a, b). Both cFcμs were polymeric when expressed in the absence of JC (Supplementary Fig. 1) and yielded structures with an average resolution of 3.9Å (Fig. 1c-d, Supplementary Fig. 2)^20^.

Cryo-EM density maps revealed that both hFcμ-tFcμ-chi3 and hFcμ-tFcμ-chi4 form pentamers, adopting near C5 symmetry with five structurally comparable Fcμ monomers arranged side-by-side in the same plane. The Fcμ monomers were separated by 72° and the C-terminal tailpieces (Tps) protruded out of the plane and formed a partially ordered barrel-like assembly with a solvent accessible tunnel (Fig. 2a, Supplementary Fig. 3a). We were able to build residues 345-559, which encompass CHμ3, CHμ4, and the loop between CH4 and Tp (termed pre-Tp loop). Structural alignment between hFcμ-tFcμ-chi3 and hFcμ-tFcμ-chi4 pentamers revealed a RMSD of 1.410 and structural alignment between each Fcμ monomer revealed a RMSD of 1.375, indicating global and local similarity between hFcμ-tFcμ-chi3 and hFcμ-tFcμ-chi4 (Supplementary Fig. 3b, c). These results signify that either the replacement of hIgM C-terminal residues 553-576 with tIgM residues 426-448 or the replacement of hIgM C-terminal residues 546-576 with tIgM residues 420-448 results in comparable structures.

**Fig. 2.**
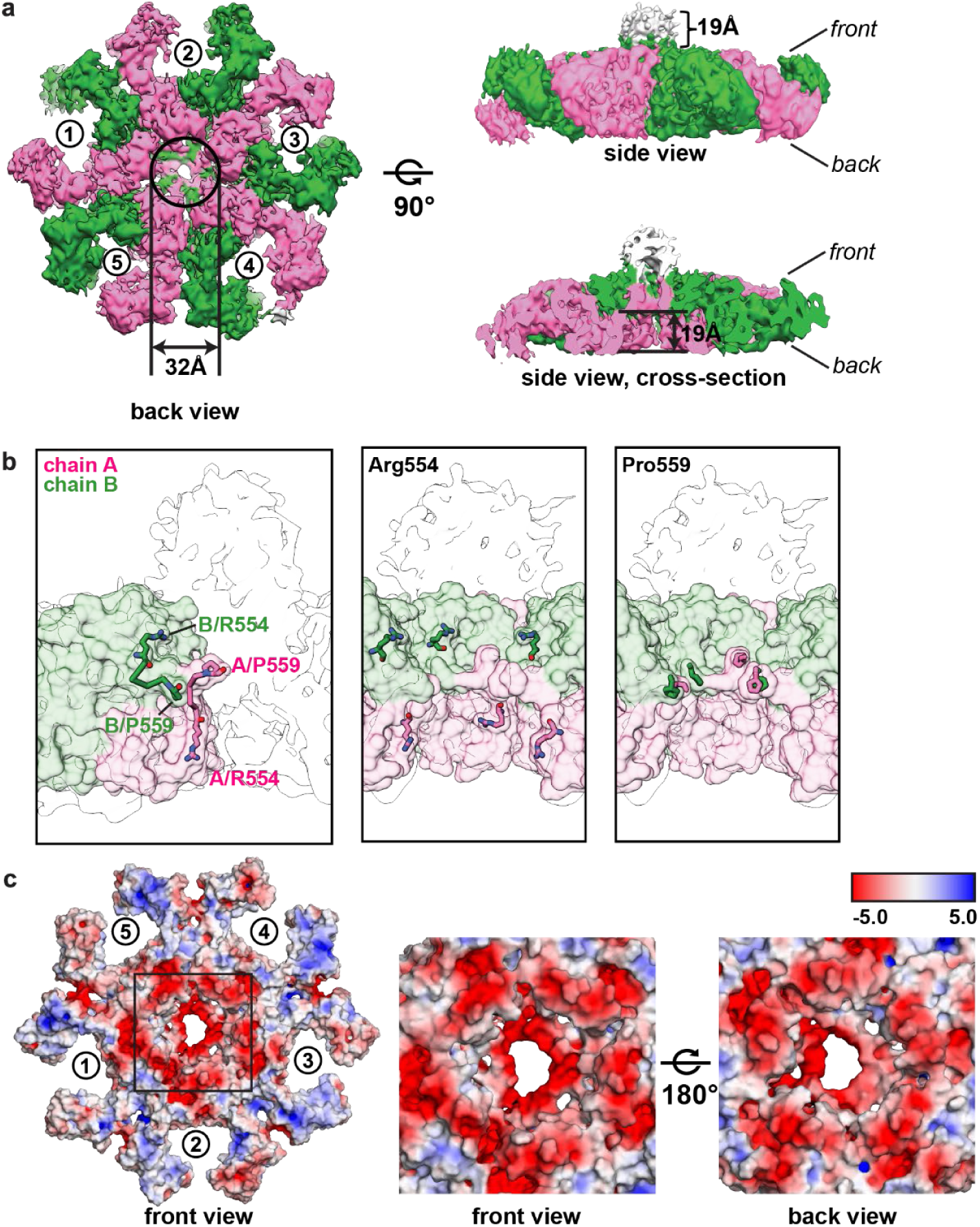
hFcμ-tFcμ-chi3 Tp assembly. **a** Cryo-EM density map of hFcμ-tFcμ-chi3 back view, side view, and side view cross-section. Density is colored and numbered as in Fig. 1 and map regions corresponding to unmodeled density are colored gray. The size of the Tp assembly protruding out of the front face, as well as the indentation on the back face are indicated. **b** *Left*, Surface representation of hFcμ-tFcμ-chi3 chains A and B with the first (Arg554) and last (Pro559) pre-Tp loop residues shown as sticks. *Middle* and *right*, Surface representation of chains E-L with Arg554 and Pro559 shown as sticks. The coloring scheme of the structure is the same as (a) and the cryo-EM density map is shown as cross-sectional tracing. **c** hFcμ-tFcμ-chi3 structure surface representation showing the electrostatic potential. The boxed region is enlarged in panels on the right.

Both hFcμ-tFcμ-chi3 and hFcμ-tFcμ-chi4 adopt pentameric structures that are distinct from published tFcμ and Fcμ-J containing structures. Among these differences is the location and structure of the Tps. The IgM heavy chain C-terminal Tps are evolutionarily conserved and have long been recognized as crucial for polymeric IgM assembly^21,22^. In the SFcμ-J structure (PDB ID: <u>6KXS</u>), which represents SIgM, as well as the tFcμ structure (PDB ID: <u>8GHZ</u>), Tps form two β-sheets that assemble into sandwich-like structures that stabilize the complexes through backbone hydrogen bonding between β-strands and hydrophobic interactions between the two sheets. In both species, the Tp assembly shares the same plane occupied by Fcμ monomers. However, in SFcμ-J the Tp assembly is located at the center of the molecule and does not directly interact with CHμ4 domains whereas in tFcμ the Tp assembly sits between Fcμ1 and Fcμ4 and directly contacts neighboring CHμ4 domains. In hFcμ-tFcμ-chi3 and hFcμ-tFcμ-chi4, we observed partially ordered Tps extending out of the Fcμ monomer plane, creating a front face with protruding Tps, and a back face with an indentation. The sub-volume attributed to Tps extends approximately 19Å out of the Fcμ plane and reveals structural details characteristic of β-strands, forming a tunnel at the center. The indentation on the back side of the Tps measured 32Å in diameter and 19Å in depth (Fig. 2a).

In both hFcμ-tFcμ-chi3 and hFcμ-tFcμ-chi4, we were able to build ten pre-Tp loops (residues 554-559), which connect CHμ4 domains and Tp density, and observe evidence that all ten Tps form a partially ordered structure; however, we could not unambiguously assign Tp sequences (residues 561-575). This likely arose from some degree of structural heterogeneity in the assembly. We observed that the two pre-Tp loops from each Fc monomer adopt two conformations that accommodate divergent positioning of CHμ4 domains with respect to the Tp assembly. Specifically, chains A, C, E, G, and K pre-Tp loops extend into the Tp assembly from the back face of the pentamer whereas chains B, D, F, H, and L pre-Tp loops descend from the top side where they form direct contacts with CHμ4 residues; this arrangement positions Pro559 from all chains on the same plane in the center of the molecule despite different starting positions of pre-Tp loops (Fig. 2b). The CHμ4-pre-Tp interface comprises CHμ4 α-helix residues Arg461, Arg467, Asn529 and pre-Tp residues Arg554, and Asn557, of chains B, D, F, H, L. Together the interface forms a hydrophilic surface surrounding the Tp protrusion (Fig. 2c). Among these residues, Arg461 participates in interchain polar contacts with the backbone and/or sidechains of pre-Tp loop residues in the neighboring chain (i.e. Arg461^Fc2D^ interacts with Asn557^Fc2C^). Arg554 sidechains on chains B, D, F, H, and L near the Tp protrusion may contact hydrophilic sidechains in Tp β-strands (Fig. 2b, Supplementary Fig. 4a). On the side opposite of the Tp protrusion, the indentation is mostly made up of hydrophilic residues from chains A, C, E, G, and K, including CH4 α-helix residues Arg461, Asn465, Asn529, and pre-Tp loop residues 554-556. Sidechains of Arg554 are exposed at the surface of the indentation and form intra-chain interactions with Asn529 of CHμ4 domain (Fig. 2c, Supplementary Fig. 4b). It is notable that copies of Arg461 and Arg554 participate in intra-chain interactions on both the front face (Tp protrusion) and back face (indentation), perhaps stabilizing the complex in two ways. To test this, we created hFcμ-tFcμ-chi3 and hFcμ-tFcμ-chi4 Arg461Met and Arg554Met single mutant expression constructs, both of which greatly reduced the pentamer to monomer ratio purified from transiently transfected cells (Supplementary Fig. 5).

Despite unassigned Tp residues following Pro559, the positions of pre-Tp residues together with map density extending outward suggest that two Tps from a single cFcμ monomer lie adjacent to each other and leave a solvent accessible tunnel in the center (Supplementary Fig. 4c). We speculate that this arrangement is stabilized by hydrophobic residues populating the Tps. Published structural and biochemical studies suggested that the evolutionarily conserved ultimate cysteine residue, Cys572, forms interchain disulfides^7,15,20^. We do not observe evidence for disulfide bonds in the Tp assembly map as reported for other polymeric Ig structures; however, we found that mutating Cys572 in hFcμ-tFcμ-chi3 or hFcμ-tFcμ-chi4 expression constructs, markedly reduced the ratio of pentamer to monomer produced by transiently transected cells, suggesting that Cys572 residues are crucial for pentameric cFcμ assembly (Supplementary Fig. 5).

### Cryo-EM structure of human hexameric IgM

The human-fish cFcμ structures revealed how JC-free pentamers can assemble; however, JC-free IgM is reportedly hexameric in humans. The dissimilarity raised the question of how JC-free hIgM hexamer resembles or differs from cFcμ pentamers. Studies using immune cell lines indicated that hexameric IgM is secreted by cells that do not express JC or fail to incorporate JC into IgM assembly although the functions of hexameric IgM remain debated^23–25^. A structural model of hexameric IgM assembly has been proposed using a combination of X-ray crystallography, NMR spectroscopy, and small angle X-ray scattering data^18^; however, high resolution hexameric hIgM structures remain unreported.

To gain insights into JC-free hexameric IgM assembly, we determined the cryo-EM structure of the human hexameric IgM-Fcμ core (hFcμ; lacking CHμ2 and Fabs) to an overall resolution of 3.94Å (Fig. 3a, Supplementary Fig. 6a-f). The refined hFcμ structure included 12 heavy chains, each comprising residues 345-559 and forming six superimposable Fcμ monomers arranged side by side in the same plane (Fig. 3b, Supplementary Fig. 6g). The structure exhibited near C6 symmetry with angles between adjacent Fcμ monomers being 59°, with the exceptions of Fcμ1-Fcμ6 and Fcμ3-Fcμ4 (62°), which are associated by a slightly larger distance (∼18Å) between CHμ4 domains at Fcμ-Fcμ interfaces compared to ∼11Å between other Fcμ copies (Fig. 3b-c). While CHμ3 and CHμ4 domains were well resolved up to residue Pro559, Tp residues 560-576 were poorly ordered and not built into the molecular model.

**Fig. 3.**
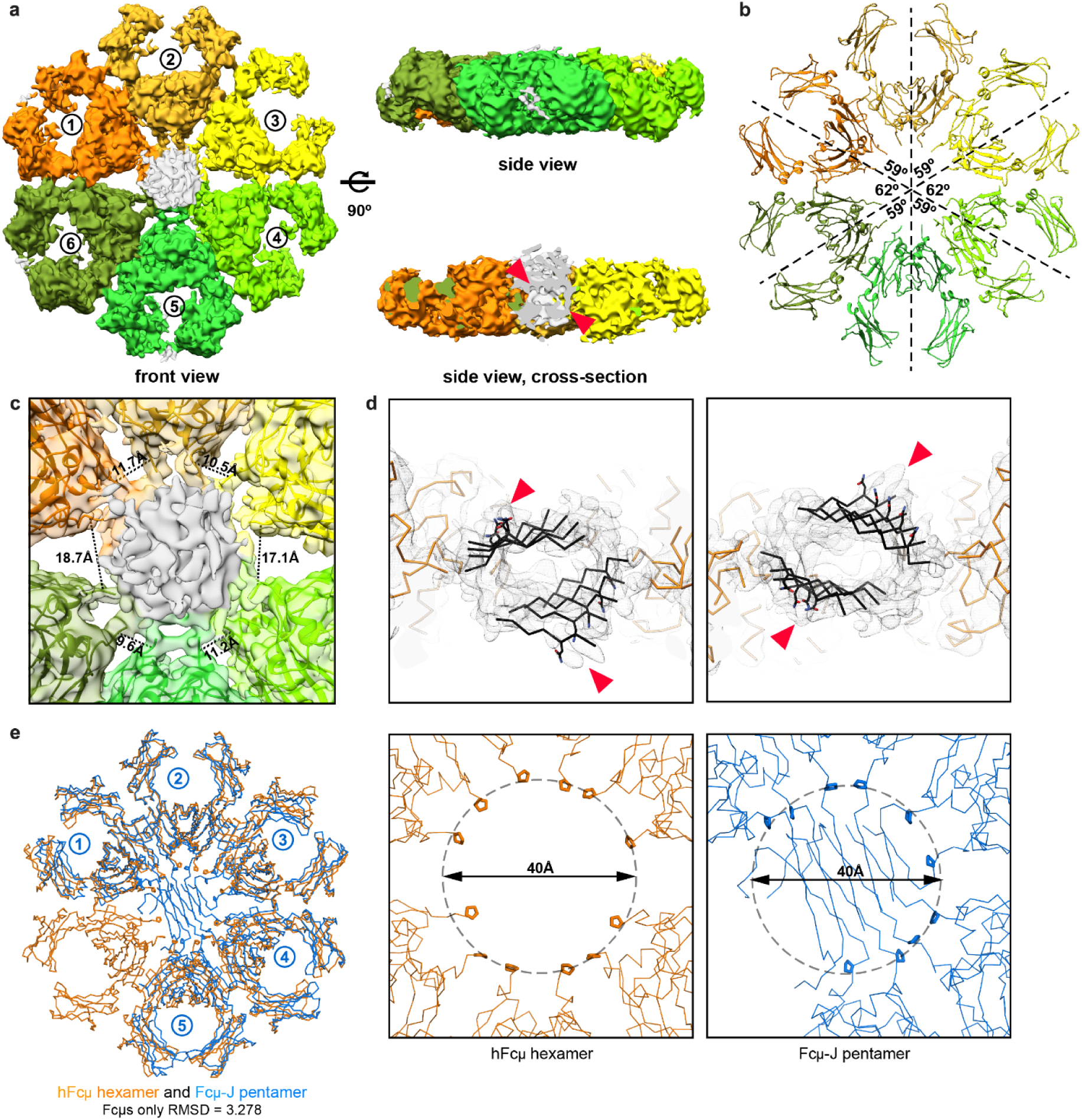
Cryo-EM map and structure of hFcμ hexamer. **a** Cryo-EM density map of hFcμ hexamer shown in three orientations with Fcμ monomers numbered and colored individually. Map regions corresponding to unmodeled density are shown in gray. **b** Cartoon representation of the hFcμ structure with angles between Fcμ monomers indicated. **c** Close-up view of the hFcμ Tp assembly with distances separating adjacent CHμ4 domains in adjacent Fcμ monomers indicated. CHμ4-CHμ4 distances were measured between Cα atoms of Lys558. Fcμ coloring scheme is the same as panel **a**. **d** Ribbon diagram showing how nine copies of Tp β-strands (*black*) fit into Tp assembly density in the context of hexameric the hFcμ structure (orange). Red triangles point to the density associated with putative glycosylation at Asn563. **e** *Left*, structural alignment between hFcμ hexamer (*orange*) and Fcμ-J pentamer (PDB ID <u>6KXS</u>, *blue*) with JC excluded. The RMSD is indicated. *Middle* and *right*, the diameter between Cα atoms of Pro559 in both structures.

Despite poor Tp resolution, the hFcμ density map showed evidence that all 12 Tps extend into the center of the hexamer to form an assembly sharing the same plane with six Fcμ monomers (Fig. 3c, Supplementary Fig. 6f). A cavity was identified in the density map at the center of the Tp sub-volume, and a β-sheet like pattern was visible in the density map when viewed at higher threshold, suggesting that the cavity identified at the center of Tp sub-volume promotes a β-sandwich-like Tp assembly that buries a hydrophobic core (Fig. 3d); however, it remains unclear how it compares to the β-sandwich formed by JC and Tps in the Fcμ-J containing pentamer structures (e.g. PDB ID: <u>6KXS</u>). The poor resolution of the map in the center of the molecule may indicate conformational flexibility in part or all of the Tp assembly (e.g. the Tps may be rigid relative to each other but flexible relative to the Fcμs) and/or the existence of multiple conformations (e.g. Tp β-strand pairing is variable) (Supplementary Fig. 6f). We were able to model two β-sheets into geometrically conceivable positions within the Tp sub-volume, one containing five poly-alanine polypeptides and the other one containing four, suggesting that the hexameric hFcμ Tp assembly can form two anti-parallel β-sheets with β-strands in each sheet lying parallel to the hFcμ plane. This hypothesis is supported by evidence for glycosylation at the conserved Asn563 site in the cryo-EM density map (Fig. 3d). Despite the absence of a refined molecular model, the Tp assembly appears to stabilize hexameric hFcμ and may promote hexamer assembly by making the process more energetically favorable (e.g. through hydrogen bonding between Tp β-strands backbones and/or formation of a hydrophobic core within the Tp assembly). The hFcμ structure also verifies that Fcμ monomers are covalently linked through disulfides between copies of Cys414 on neighboring Fcμs, as reported for pentameric Fcμ-J (Supplementary Fig. 6h)^15^.

Our hexameric IgM is planar like Fcμ-J pentamer, differing from the previously reported multi-method model of hexameric IgM, which adopted a mushroom-like shape with CHμ3 and CHμ4 domains lying on different planes^18^. Global alignment between hFcμ (hexamer) and Fcμ-J (pentamer from <u>6KXS</u>, JC excluded in the alignment) reveals that structures are nearly superimposable with highly comparable Fcμ-Fcμ angles, excluding the 123° gap between Fcμ1 and Fcμ5 in Fcμ-J, which is occupied by JC and serves as the binding site for secretory component (Fig. 3e, Supplementary Fig. 7a); in the hFcμ hexamer, this space is occupied by an additional Fcμ monomer instead. We found that the central volume to accommodate Tps was similar in hFcμ hexamer and Fcμ-J pentamer, as evidenced by a common diameter of ∼40Å between N-termini of Tp β-strands (measured between copies of Pro559) (Fig. 3e). Our analysis also revealed features that may impact the positions of CHμ2 domains and Fabs compared to several Fcμ-J containing structures, which reportedly adopt distinct Fcμ1 monomer positions. Specifically, the asymmetrical binding of JC to Fcμ1 in Fcμ-J pentamer confers CHμ3 domain tilting and limited CHμ2 flexibility, both expected to impact the positions Fcμ1 monomer Fabs^17^. In hexameric hFcμ, we do not observe marked CHμ3 domain tilting; rather, the six Fcμ monomers exhibited high structural similarity and aligned well with Fcμ2-5 in Fcμ-J structures, but deviated from Fcμ1^15,17^ (Supplementary Fig. 6g, Supplementary Fig. 7b).

### Comparison of cFcμ pentamers to human IgM

The cFcμ pentamers and hFcμ hexamer structures reported here suggest alternative modes of polymeric Ig assembly in the absence of JC. To better understand the structural differences between cFcμ pentamers and hIgM polymeric species, we conducted global and local alignment between hFcμ-tFcμ-chi3, Fcμ-J pentamer, and hFcμ hexamer. While individual Fcμ monomers are relatively comparable, hFcμ-tFcμ-chi3 was not globally superimposable with either Fcμ-J pentamer or hFcμ hexamer (Fig. 4a, Supplementary Fig. 8a). A primary difference is distinct geometrical relationships between each Fcμ and associated domains. Adjacent Fcμ monomers in hIgM adopt angles of ∼60° both in the presence and absence of JC (Fig. 3b, Supplementary Fig. 7a) whereas cFcμ pentamers adopt a larger Fcμ-Fcμ angle of about 72° (Supplementary Fig. 3a); this results in cFcμ structures having a smaller diameter of 166Å than hIgM structures, which have a diameter of ∼180Å (Supplementary Fig. 7c). This implies that the distances from the center of the molecule to antigen binding regions are conserved in hexameric and pentameric hIgM but are different in our engineered cFcμs. The Fcμ arrangement in hIgM also results in a larger diameter of ∼40Å between the C-termini of CHμ4 domains (measured between copies of Pro559) to accommodate the Tp assembly compared to cFcμs, where ten copies of Pro559 are arranged in a smaller circle with a diameter of ∼20Å that likely constrains the possible positions of the Tps and prevents a β-sandwich Tp assembly from sharing the same plane as the Fcμ monomers as observed in hIgM structures (Fig. 3e, Fig. 4b). This might account for the unique Tp assembly protrusion we observe in the cFcμs. It is also conceivable that differences between structures arise in part from differences in glycosylation of Tp residues. For example, in cFcμ structures the protruding Tp assembly and its proximity to CHμ4 domains may result from the absence of ordered glycosylation at Asn563, a conserved potential N-linked glycosylation site in both tIgM and hIgM sequences (Supplementary Fig. 9). While this site is indeed glycosylated in both Fcμ-J pentamer and hFcμ hexamer structures, the equivalent site in tIgM (Asn436) does not exhibit evidence for glycosylation suggesting that the tIgM sequences and the structure it adopts in both tFcμ and cFcμs do not support ordered glycosylation^15,20^.

**Fig. 4.**
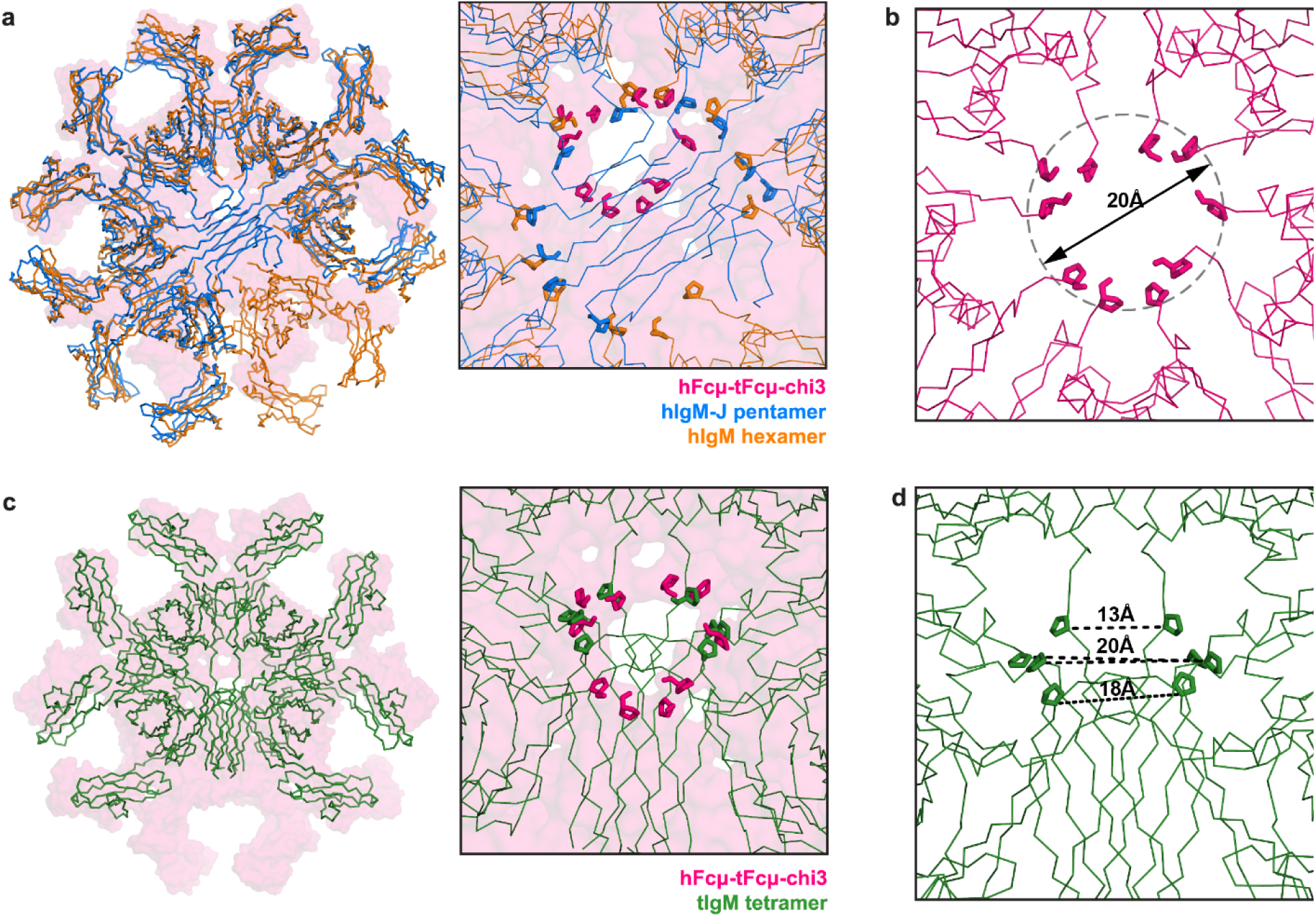
Comparison between hFcμ-tFcμ-chi3, Fcμ-J pentamer, hFcμ hexamer, and tFcμ tetramer. **a** Left, structural alignment between hFcμ-tFcμ-chi3 pentamer (pink molecular surface), Fcμ-J pentamer (JC removed; blue ribbon), and hFcμ hexamer (orange ribbon). *Right*, enlarged view of the Tp assembly with Pro559 shown as sticks. **b** Close-up of hFcμ-tFcμ-chi3 Tp assembly (pink ribbon) with Pro559 shown as sticks. **c** *Left*, structural alignment between hFcμ-tFcμ-chi3 pentamer (pink molecular surface) and tFcμ tetramer (green ribbon). *Right*, Enlarged view of the Tp assembly with Pro559 shown as sticks. **d** Enlarged view of the tFcμ tetramer Tp assembly (green ribbon) with Pro559 shown as sticks and distances labeled.

Distinct geometrical relationships between Fcμs in cFcμs and hIgM structures are also associated with differences in the separation between CHμ3 domains within each Fcμ monomer, as well as the CHμ4-CHμ4 contact surface between adjacent Fcμs. For example, within each cFcμ, two CHμ3 domains are separated by 24.6-28.6Å (measured between Cα atoms of residue Ile345), whereas the distance reduces to 18.8-22.4Å in hFcμ hexamer and 18.7-21.4Å in Fcμ-J pentamer (Supplementary Fig. 7c). The more compact geometry of cFcμ pentamers is correlated with a slightly larger CHμ4-CHμ4 contact surface between adjacent Fcs, encompassing an average of 495Å^2^ surface area, compared to an average of 401Å^2^ in the hIgM-J pentamer. This was unexpected since residues comprising the CHμ4-CHμ4 interface, which encompass CHμ4 domain β-strands C, F, and G plus loops flanking these strands, are either identical or evolutionarily conserved between hIgM and cFcμs (see sequence alignment in Fig. 1b); however we observe some differences in contacts in the loop between β-strands C and D, and the loop between β-strands F and G (Supplementary Fig. 7d). In cFcμs these two loops are direct towards the core of the CHμ4-CHμ4 interface and form a larger contact area. This observation suggests malleability in Fcμ-Fcμ interfaces.

Despite noted differences among cFcμ and hIgM structures, we find that the evolutionarily conserved Cys414 residue, located in the α2 helix of CHμ3 domain, does form interchain disulfides in all four structures (Supplementary Fig. 3d, Supplementary Fig. 6h). Mutation of Cys414 reduced the pentamer:monomer ratio of recombinantly expressed cFcμ proteins from 68% to 14% in hFcμ-tFcμ-chi3, and from 58% to 16% in hFcμ-tFcμ-chi4 (Supplementary Fig. 5). These results are consistent with previous biochemical studies conducted on recombinantly expressed hIgM, in which mutation of Cys414 reduced the percentage of polymeric complexes^7^. Intriguingly, this cysteine is not conserved in lower vertebrate species (i.e. cartilaginous fish and teleosts) and therefore does not contribute to interchain interactions in the tetrameric tFcμ structure (Supplementary Fig. 9)^20^.

In hIgM, each Fcμ promotes antibody effector functions activation by binding to Fc receptors. The structural basis of Fcμ recognition by IgM-specific Fc receptor, FcμR/TOSO, was revealed in published cryo-EM structures, which show that each Fcμ monomer provide up to two FcμR binding sites, located on each side of the CHμ4-CHμ4 homodimer. In Fcμ-J (lacking SC), eight FcμR binding sites are available on Fcμ2-5; this allows simultaneous binding of up to eight copies of FcμR. However in SFcμ-J, FcμR binding to the SC-bound face was abolished, eliminating four sites^26,27^. In our cFcμ structures, FcμR binding sites on CHμ4 domains are solvent accessible; however modeling suggests that only up to four copies of FcμR can be docked simultaneously onto cFcμ without encountering steric clashes.

### Comparison of cFcμ pentamers to tFcμ tetramer structure

While tFcμ tetramer and cFcμ pentamers adopt JC-free structures, they are not superimposable and do not share the same Tp assembly location (Fig. 4c). tIgM tetramer adopts a C2 symmetry, with the Tp assembly situated between Fcμ1 and Fcμ4 and does not directly contact Fcμ2 or Fcμ3. In contrast, cFcμs adopt C5 symmetry and the Tp assembly is nested at the center of the molecule, contacting all five Fc monomers. The location of Tp assembly differs between tFcμ and cFcμs, leading to distinct spatial distribution of the last residue before Tp β-strands, Pro432 in tFcμ and Pro559 in cFcμs. In cFcμs ten copies of Pro559 occupy the roughly the same plane, whereas in tFcμ tetramer, eight copies of Pro432 are distributed across three planes due to asymmetric positioning of the Tp assembly. The distance between Pro432 on opposing chains varies between 13Å and 20Å, comparable to the diameter of Pro559 residues in cFcμs which measures at ∼20Å but much smaller than the hIgM structures (Fig. 4b-d). Furthermore, in cFcμ, Tps from the same cFcμ monomer appear to run side-by-side to form a Tp assembly, resembling Fcμ-J pentamer and possibly, the hFcμ hexamer (Supplementary Fig. 4c-d), which is distinct from tFcμ tetramer where Tps from each tFcμ monomer are separated by one Tp from the adjacent tFcμ monomer, creating an alternating pattern in Tp β-sheets (Supplementary Fig. 8b).

In both hFcμ-tFcμ-chi3 and hFcμ-tFcμ-chi4, pre-Tp loops and Tps contain solely tIgM residues that share a up to 40% sequence similarity with hIgM (Supplementary Fig. 9), yet these residues form alternative structures. Our prior work revealed tFcμ pre-Tp loops adopting multiple conformations and mediating interactions between the Tp assembly and CHμ4 domains^20^. For example, in tFcμ two heavy chain pre-Tp loops (A and H) neighboring the Tp assembly adopt a unique fold (“A-type” fold) and wrap around the N-termini of Tp β-sheets, positioning pre-Tp loop residue Arg427 to interact with Tp residue Ser438 on other chains (Supplementary Fig. 8b). Structural alignments between individual cFcμ monomers and tFcμ monomers revealed no evidence for the “A-type” fold even though residues forming pre-Tp loops are present in both cFcμs (Supplementary Fig. 8a). Nevertheless, interactions between pre-Tp loops and the Tp assembly are still sterically possible in cFcμ pentamers given that pre-Tp loops of chains B, D, F, H, and L constitute the CHμ4-Tp interface surrounding the Tp assembly. Arg554, equivalent to tIgM residue Arg427, is positioned to interact with Tp residues in a chain-specific fashion, though specific interactions could not be determined due to limited resolution in the Tp assembly (Supplementary Fig. 4a). Notably, this interaction is absent in hIgM and copies of the equivalent residue, Lys554, are located distal from Tp strand residues.

### Fusion of sdAb targeting CDI to cFcμs produce toxin neutralizing polymeric antibodies

Having characterized the structures of cFcμs and found differences compared to human and teleost counterparts, we sought to determine if the structures could serve as engineered, polymeric antibodies, providing a novel JC-free scaffold with unique structural characteristics. To investigate the compatibility of cFcμ scaffolds with single domain antibody (sdAb) fusions, we created an expression construct that linked previously characterized sdAb A20.1 to the N-termini of both hFcμ-tFcμ-chi3 and hFcμ-tFcμ-chi4, named sdA20.1-cFcμs (Fig. 5a). The sdA20.1 targets the opportunistic human pathogen, *Clostridioides difficile* (*C.difficile*) toxin TcdA^28^. TcdA an important factor in the pathogenesis of *C. difficile* infection (CDI), binding to host epithelial cells surface receptors and causing cytotoxicity associated with barrier disruption^29^. Limiting toxin activity through administration of toxin-binding monoclonal IgGs is a proven therapeutic approach for treatment of recurrent CDI^30^.

**Fig. 5.**
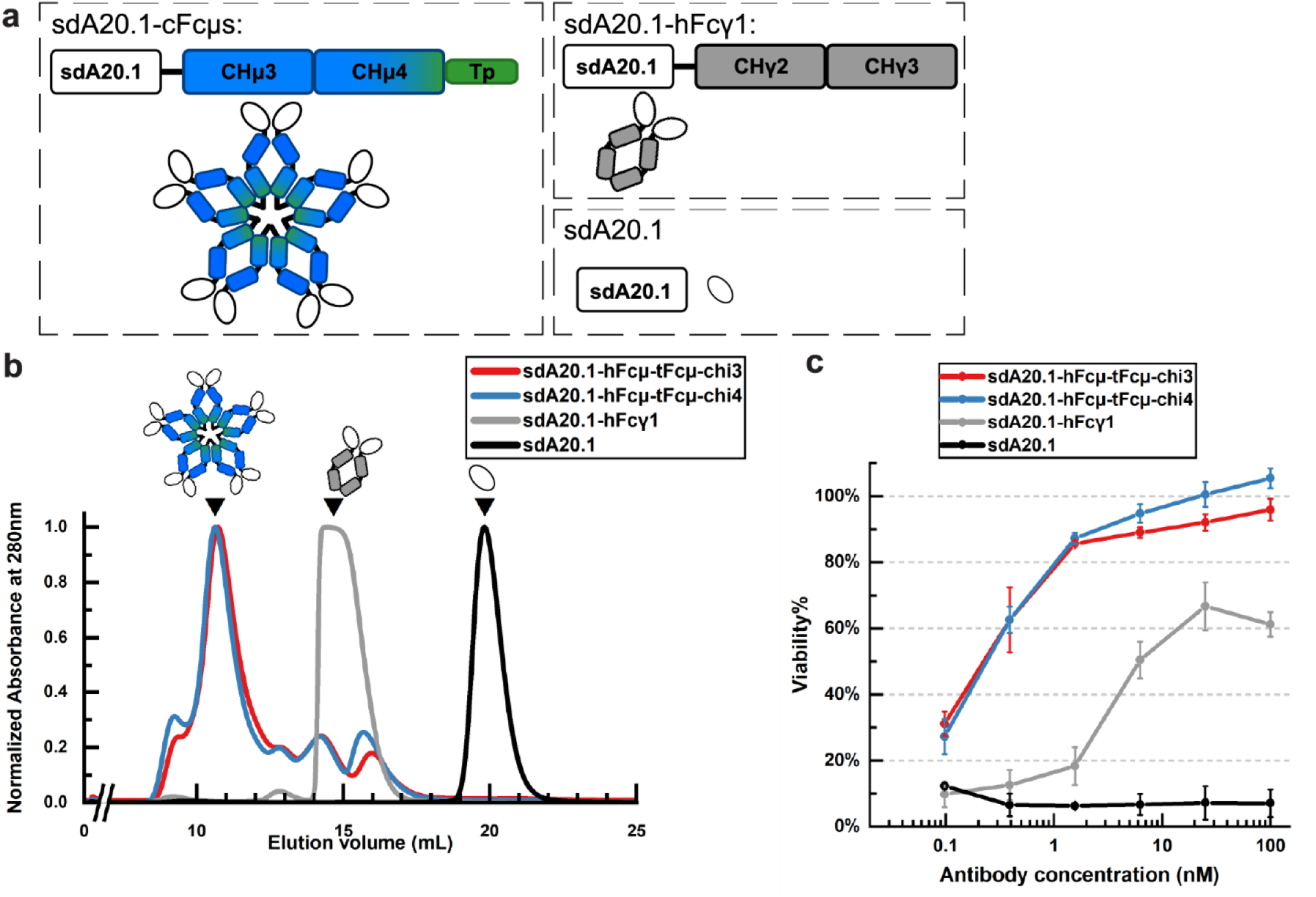
sdA20.1-cFcμs and *C. difficile* TcdA neutralization. **a** Schematic of pentameric sdA20.1-cFcμs, sdA20.1-hFcγ1, and sdA20.1. **b** Size exclusion column chromatograms of purified, recombinantly expressed sdA20.1-cFcμs, sdA20.1-hFcγ1, and sdA20.1. Triangles indicate the peaks collected for subsequent Vero cell-based cytotoxicity neutralization assays against TcdA. **c** Concentration-dependent neutralization of TcdA (50pM) by the indicated sdA20.1-containing antibodies. Data shown are the result of triplicate measurements.

The sdA20.1-cFcμ constructs were recombinantly expressed in human cell lines without JC, producing heterogeneous antibodies with variable polymeric states ranging from monomer (8-15%) to pentamer (62-69%) (Fig. 5a, b). To determine if pentameric sdA20.1-cFcμs retain the binding activity and potency of sdA20.1, we conducted Vero cell-based cytotoxicity neutralization assays against TcdA. Vero cell cultures were incubated with mixture of 50pM TcdA and varying concentrations of sdA20.1-cFcμs, or controls sdA20.1, and sdA20.1 fused to the Fc of human IgG1 (sdA20.1-hFcγ1). Following a 68-72 hour incubation, cell viability was measured using an Alamar Blue assay (Fig. 5a)^31,32^. Whereas 50pM TcdA alone caused ∼100% Vero-cell death in the absence of antibodies, both sdA20.1-cFcμs provided protection against TcdA toxicity, exhibiting TcdA neutralization at concentrations as low as 0.1nM and fully restored cell viability at 100nM (Fig. 5c). Notably, pentameric sdA20.1-cFcμs showed higher potency than sdA20.1-hFcγ1 and sdA20.1, the former recovering up to ∼60% cell viability at 100nM concentration and the latter not showing significant protection from TcdA. Together data indicate that pentameric sdA20.1-cFcμs can outperform concentration matched monomeric species suggesting that hFcμ linked to other sdAbs may provide neutralization advantages against other antigens.

## DISCUSSION

The polymeric (p) and secretory (S) Igs are large multi-protein complexes that connect rigid Fc cores of varying size, shape and composition to flexible Fabs, thereby providing the host with unique ways to bind antigen and unique effector mechanisms to counter pathogens and promote microbial homeostasis. These characteristics require intricate molecular assembly mechanisms that until recently, remained poorly understood. While our understanding of the structural mechanisms underlying JC-dependent pIg assembly has advanced in recent years, knowledge of JC-free assembly has remained limited^15–17,33,34^. Our previously reported tFcμ structure (PDB ID: <u>8GHZ</u>) revealed that teleost, JC-free assembly incorporates distinctive tFcµ folding patterns not observed in Fcμ-J pentamer-containing structures or pIgA structures. This culminates in a Tp assembly situated between two Fcμs rather than the center of the molecule like Fcμ-J, suggesting potentially divergent biophysical mechanisms underlying JC-free versus JC-dependent assembly. Yet, accompanying biochemical characterization of cFcμs containing both tIgM and hIgM motifs demonstrated that tFcµ C-terminal residues can facilitate JC-free pIg assembly when fused to hIgM heavy chain, indicating mechanistic interchangeability between pIg-forming heavy chain sequences and their assembly mechanisms^20^. Here, our cFcμ structures have revealed that pIg assembly mechanism plasticity extends beyond interchangeability among naturally selected structures and can culminate in the unique structures observed for hFcμ-tFcμ-chi3 and hFcμ-tFcμ-chi4.

We can pinpoint the minimum requirements for building cFcμ’s unique structures to include the hIgM heavy chain residues 343-553 and tIgM Tp residues 426-448 (see hFcμ-tFcμ-chi3 sequence, Fig. 1a). It is notable that hFcμ-tFcμ-chi3 and hFcμ-tFcμ-chi4 are nearly superimposable despite additional teleost residues (residues 547-553) upstream of the Tp in hFcμ-tFcμ-chi3. We had anticipated differences associated with the inclusion of these residues in hFcμ-tFcμ-chi3 because in the tFcμ structure, they adopt different positions and β-strand pairing in two of the eight heavy chains (those with the so called “A-type” fold)^20^. However, in hFcμ-tFcμ-chi3 they occupy the same positions as equivalent residues from the human IgM sequence and thus do not promote an alternative folding pattern or alternating β-strand pairing (Supplementary Fig. 3b, c, Supplementary Fig. 8a).

Given the conservation between the teleost and human IgM Tps (40% sequence similarity) and the observation that these Tps are associated with tetrameric tIgM or hexameric hIgM, respectively, in their cognate structures, our data raise the question of why cFcμs adopt a pentameric structure. In addition to Tp β-sheets, other hIgM heavy chain motifs reportedly contribute to pIgM assembly, including inter-Fc disulfides between copies of CHμ3 Cys414 as well as interchain disulfides between copies of Tp Cys575. While Cys414 is absent from the tIgM sequence, a C-terminal cysteine is conserved in all jawed-vertebrates, though its position varies. In most species it is the penultimate residue but is fourth from the C-terminus in teleosts. In the case of tIgM tetramer, the Fc-Fc interfaces lack disulfides and the Tp cysteine residue Cys445 is incompletely oxidized and therefore likely forms transient inter-chain disulfides, as evidenced by both biochemical assays and the high-resolution cryo-EM structure^20,22^. The incorporation of C-terminal residues in cFcμs does not affect inter-Fc disulfides between Cys414 but shifted the position of penultimate Cys575 in hIgM Tp to Cys572 preceding three C-terminal residues. The disulfides formed by Cys414 are associated with a closer distance between CHμ3s of adjacent monomers and likely favor C5 symmetry over C2 symmetry, resulting in an elbow-to-elbow Fc-Fc interface in cFcμs pentamers. In contrast, cFcμs Cys572 adopts a distinct position further away from the C-terminus. We speculate that this position may impact interactions with ERp44, a chaperone essential to endogenous hIgM assembly that facilitates inter-monomer disulfides between copies of Cys575^36^, possibly resulting in incomplete disulfide formation between copies of Cys572. This hypothesis is consistent with the minimal population (∼10%) of polymeric cFcμ in Cys572Ala SEC-MALS data, which is otherwise populated by monomers (Supplementary Fig. 5). Together, the combination of covalently linked CHμ3s and Cys572’s distinct postion may provide variable driving forces that tip the cFcμ assembly process toward pentamer formation.

The cryo-EM structure of hFcμ hexamer highlights the plasticity of pIg assembly mechanisms in the context of a naturally occurring, JC-free process that results in functionally relevant similarities and differences in structure compared to pentameric Fcμ-J^1^. hFcμ hexamer adopts a hexagonal geometry and is superimposable with hFcμ-J pentamer except for the sixth Fcμ monomer occupying the gap observed in Fcμ-J pentamer. In hexameric IgM, elbow-to-elbow contacts between all six Fcμ monomers may restrict tilting of CH3 domains and increase the rigidity of the IgM’s core structure; however, our data suggest that within IgM hexamer the Tp assembly may have more flexibility relative to the covalently linked Fcμs when compared to the equivalent of Fcμ-J pentamer, as evidenced by our inability to unambiguously assign residues to the Tp assembly sub-volume.

The observation that pentameric cFcμ and hexameric hFcμ Tp assemblies adopt distinct structures from JC-containing pIgs (e.g. Fcμ-J) supports the previously proposed JC-templating model for Tp assembly. The JC-templating model states that JC adopts an anti-parallel micro β-sandwich core that serves as a template for Tp β-strand incorporation into two β-sheets constituting the Tp assembly^33,37^. This model is supported by several examples of JC-containing pIg structures including JC-containing human SIgM core (PDB ID: <u>6KXS</u>) and SIgA (PDB ID: <u>6UE7</u>). In these two structures, the anti-parallel micro β-sandwich of JC establishes the opposing directions of β-sheets; two Tps of the same Fc monomer are incorporated into the Tp assembly in a pair-wise fashion by extending one face of the anti-parallel β-sheets. In SIgM, this occurs until the addition of the fifth Fc monomer, whose Tps add to each side of β-sandwich. The Tp assembly is essentially capped by two JC β-strands on the front side and by one JC β-strand on the back side^15,33^. In contrast, cFcμ and hFcμ structures lacking JC reveal Tp assemblies that deviate from the canonical JC-templated Tp assembly. The cFcμs adopt tunnel-like structures (Supplementary Fig. 4c) and hFcμ hexamer adopts one or more β-sheet-like structures with higher apparent flexibility than Fcμ-J. Despite this, the map showed evidence for β-sheets similar to Fcμ-J pentamer suggesting that Tp β-strand interactions occur but are transient, and/or can adopt multiple positions and/or adopt an alternating pattern of Tp β-strand pairing. Any of these possibilities would suggest that in JC-containing structures, JC constrains the order and the final structure of Tp assembly as the templating model suggests. Alternatively, the hFcμ hexamer Tp assembly may adopt rigid β-sheets similar to Fcμ-J pentamer that are flexible relative to the linked Fcμs that encircle it (i.e. a floating disk at the center), a possibility that could render the JC unnecessary for templating the β-sheet structures similar to those observed in Fcμ-J, but critical for ensuring that less than six Fcμ monomers are added, for stabilizing the complex, and for providing pIgR (and SC) binding sites.

Our ability to produce cFcμ with unique structures not only provides insight into the plasticity and driving factors behind IgM assembly but also provides a novel approach to engineering IgMs. Published high-profile reports include the development of avidity-enhanced human IgM-linked sdAb platforms that protect against SARS-Cov-2 infection in animal models when delivered intravenously^38^. Here we show that cFcμ can be similarly linked to sdAbs (e.g. sdA20.1-cFcμs) and neutralize toxins secreted by human pathogen, *C. difficile*. While future efforts will be needed to optimize this platform to target antigens, the approach provides a scaffold with geometry that almost certainly differs from existing platforms based on human pentameric or hexameric IgM. We speculate that this differing geometry may provide functional advantages, providing different spacing between antigen binding sites and perhaps modulate the ability to control effector functions.

## METHODS

### Construct design and protein expression of Fcs

The amino acid sequence of *Oncorhynchus mykiss* IgM heavy chain secretory form (GenBank: <u>AAW66974.1</u>) and *Homo sapiens* IgM heavy chain constant domains (UniProtKB: <u>P01871.5</u>) were obtained from the NCBI database (https://www.ncbi.nlm.nih.gov/). DNA sequences encoding a signal peptide (MDAMKRGLCCVLLLCGAVFVSPAGA), a hexa-histidine tag, and the hFcµ, tFcµ, or hFcµ-tFcµ-cFc (cFcµ) coding sequence were codon optimized for human cell expression, synthesized (Integrated DNA Technologies, Inc.), and cloned into mammalian transient expression vector pD2610-v1 (Atum). Design and sequence of hFcµ-tFcµ-cFc constructs used in this study have been previously described in Lyu et al.^20^. Resulting plasmid DNA was transiently transfected into HEK Expi293F cell line (Gibco) using the ExpiFectamine 293 Transfection Kit (Gibco). Six days after transfection, supernatants were harvested and His-tagged proteins were purified by affinity chromatography using Ni-NTA Agarose (Qiagen), exchanged into 1x TBS buffer (20mM Tris-HCl and 150mL NaCl, pH 7.8), and then subjected to size exclusion chromatography (SEC) on a Superose 6 Increase 10/300 column (Cytiva) installed on an ÄKTA pure FPLC system (Cytiva). Single mutant cFcµ variants were expressed and purified in the same way.

### Construct design and protein expression of sdAb-cFcµ antibodies

The amino acid sequence of single domain antibody A20.1 was obtained from Murase et al.^39^, reverse translated into nucleotide sequences, and codon optimized for human cell expression. Nucleotide constructs encoding sdAb, linker, and Fcµ (termed sdAb-cFcµ) were cloned into mammalian transient expression vector pD2610-v1 (Atum). Resulting plasmids were transiently transfected as described above. sdAb-cFcμ proteins were purified by affinity chromatography using POROS CaptureSelect IgM-XL Affinity Matrix (Thermo Fisher) or MabSelect antibody purification chromatography resin (Cytiva), exchanged into 1x PBS buffer (137mM NaCl, 2.7mM KCl, 10mM Na_2_HPO_4_, 1.8mM KH_2_PO_4_, pH 7.4), and subjected to SEC on a Superdex 200 Increase 10/300 column (Cytiva) installed on an ÄKTA pure FPLC system.

### SEC-Multiangle light scattering (SEC-MALS) analyses

Recombinant protein samples were expressed and purified as described above. After purification by SEC, fractions in each peak were pooled, concentrated to ∼2g/L, filtered, and subjected to SEC-MALS analysis. For each protein, 100uL injection was applied onto a Superose 6 Increase 10/300 column (Cytiva) installed on NGC Discover 10 Chromatography System (Bio-rad) with a flow rate of 0.75 mL/min at 25°C. The scattered light intensity of the eluant was recorded using a DAWN-HELEOS MALS detector (Wyatt Technology) at 18 different angles operating at 658 nm and the index of refraction (RI) was collected by an OPTILAB refractometer (Wyatt Technology). MALS and RI raw data were analyzed using the ASTRA software (Wyatt Technology) after calibration of MALS detectors were performed using commercial bovine serum albumin (BSA). Weight average molar mass of each data slice was determined through construction of a Debye plot (Rθ/(K*c) against sin^2^(θ/2) plot) and polynomial fitting in such plot. The overall average molecular weight of species under a peak was the weight-average of the determined weight average molar mass.

### Cryo-EM grid preparation, data collection

Post SEC, the highest molecular weight peaks were pooled from hFcµ-tFcµ-chi3, hFcµ-tFcµ-chi4, and hFcµ preparations. Proteins were diluted to 0.2 g/L in 1x TBS buffer and filtered prior to grid freezing. Quantifoil R 2/2 UT 400 Mesh Copper grids (Quantifoil) were glow discharged at 15mA current for 90 seconds. Cryo-EM grids were prepared at 4°C, 100% humidity, with blot force of 3, wait time of 2 seconds, and blotting time of 2 seconds in a Vitrobot Mark IV (Thermo Fisher). Movies were collected on Titan Krios G4 Cryo-TEM (Thermo Fisher) operating at 300kV and movies were recorded using SerialEM on a post-GIF K3 Direct Electron Detector (Gatan). 1,986 untilted movies of hFcµ-tFcµ-chi3 pentamer, 2,068 untilted movies of hFcµ-tFcµ-chi4 pentamer, and 2,224 untilted movies of hFcµ hexamer were collected in super resolution mode with a corrected pixel size of 0.529Å/pix, and 57.35 e/Å^2^ total dose. 2,026 tilted movies of hFcµ-tFcµ-chi3 pentamer and 2,043 tilted movies of hFcµ-tFcµ-chi4 pentamer were collected on a 30° tilted stage under the same parameters.

### Cryo-EM data processing

Untilted movies were imported into cryoSPARC^40^ and patch motion corrected with a binning factor of 2. Micrographs were then subjected to patch-based CTF estimation and then manually curated for average defocus, CTF fit, ice thickness, etc. 1,549 micrographs were accepted for hFcµ-tFcµ-chi3, 1,679 micrographs were accepted for hFcµ-tFcµ-chi4, and 1,594 micrographs were accepted for hFcµ hexamer. Tilted movies were also imported and pre-processed in the same manner as untilted movies; 1,703 tilted micrographs were accepted for hFcµ-tFcµ-chi3 and 1,855 tilted micrographs were accepted for hFcµ-tFcµ-chi4.

Several reiterations of particle picking, particle extraction, and 2D classification were first performed on untilted micrographs and then particles in selected 2D classes were re-extracted from micrographs to improve data quality. From the hFcµ hexamer dataset, an initial stack of 369,073 particles was used for ab initio reconstruction into three classes and subsequently heterogeneous refinement. The largest class, containing 216,538 particles, was used in non-uniform refinement^41^ under C1 symmetry and local refinements, resulting in a map of 3.93Å overall resolution. Local resolution of density map was evaluated in cryoSPARC.

For cFcµs, good 2D classes selected from untilted micrographs were used as templates to pick particles from titled micrographs, which underwent reiterations of 2D classification and selection. Particles selected from both untilted and tilted micrographs were combined to generate a stack of 528,158 particles from hFcµ-tFcµ-chi3 micrographs, and a stack of 477,625 particles from hFcµ-tFcµ-chi4 micrographs. Ab initio reconstruction and subsequent heterogeneous refinement^41^ were run on each particle stack into three classes. The best class was used for non-uniform refinement under C1 symmetry and local refinements, resulting a hFcµ-tFcµ-chi3 map of 3.87Å overall resolution and a hFcµ-tFcµ-chi4 map of 3.92Å overall resolution. Local resolution of density maps was evaluated in cryoSPARC.

### Molecular model building, refinement, and validation

Homology models of hFcµ-tFcµ-chi3 monomer and hFcµ-tFcµ-chi4 monomer were generated using SWISS-MODEL^42^ using hFcµ1 (Tp excluded) from a Fcµ-J pentamer (PDB ID:6KSX) as a template. For each cFcµ, five copies of monomer homology model were docked into their respective real-space density maps using UCSF Chimera (developed by the Resource for Biocomputing, Visualization, and Informatics at the University of California, San Francisco, with support from NIH P41-GM103311)^43^. After initial docking, all CHµ3 and CHµ4 domains were re-docked individually to improve fitting; CHµ3 and CHµ4 were re-connected in *Coot* Molecular Graphics Package^44^. The resulting models of pentameric cFcµs were subjected to reiterations of real-space refinement in PHENIX^45^ and manual adjustments in *Coot*.

Six copies of hFcµ1 (Tp excluded) from Fcµ-J pentamer (PDB ID: <u>6KXS</u>) were docked into real-space density map of JC-free hIgM hexamer using UCSF Chimera. After initial docking, individual CHµ3 and CHµ4 domains were re-docked individually and then reconnected in *Coot*. The result hFcµ hexamer model was subjected to reiterations pf automatic real-space refinement in PHENIX^45^ and manual adjustments in *Coot*^44^.The final molecular models of hFcµ-tFcµ-chi3, hFcµ-tFcµ-chi4, and hIgM were evaluated by Phenix EM Validation, MolProbity, and EMRinger^45,46^. Data collection and refinement statistics are provided in Supplementary Table 1.

### Sequence alignments and structural alignments

Sequence similarity scores for short sequences (Tps) were calculated using https://www.bioinformatics.org/sms2/ident_sim.html. Sequence alignments between tFcμ, hFcμ, and cFcµs were made using ClustalOmega and alignment figures were made using Espript3^47,48^. Structural alignments were performed using the “align” function in PyMOL^49^. Tps were removed for structural alignment between tFcμ, hFcμ, and cFcµ monomers, but retained for global structural alignment of indicated structures unless specified otherwise in figure legend.

### Density map and molecular model measurements

For measurements of cryo-EM density maps, volume tracers were placed directly on the density maps and the distances between volume tracers were measured using UCSF Chimera. The height of the cFcµ Tp protrusion and depth of cFcµ back-face indentation were reported as averages of at least three measurements. To measure Fcµ-Fcµ angles UCSF Chimera centroids were defined for each individual Fcµ monomer (with Tps removed) without mass weighting. The centroid of pIg structures was defined without mass weighting. Then, centroids of individual Fcµs were each connected to the centroid of the pIg. The connecting lines were considered as axes for each Fcµ monomer and Fcµ-Fcµ angles were measured as the angles between these axes. Distance measurements were performed in PyMOL between Cα atoms of residues of interest.

### Cytotoxicity assay using Vero cells

Vero cells (ATCC-CCL-81) were maintained at 8% CO_2_ at 37°C. On day 0, Vero cells were seeded into tissue culture surface treated 96-well plates at 10^4^ cells/well in complete media made up of Dulbecco’s Modified Eagle Medium (DMEM, Gibco) and 10% Fetal Bovine Serum (FBS, Gibco). Experimental antibodies (e.g. sdA20.1-cFcμs) were diluted to create a gradient and were pre-incubated with 50pM commercial *Clostridioides difficile* toxin TcdA (Sigma-Aldrich) at 8% CO_2_ at 37°C for one hour. Medium only wells and TcdA only wells were also incubated as negative controls and toxin only controls, respectively. Pre-incubated mixtures of antibodies and toxin were added to Vero cells 20-24 hours post seeding (day 1). 68-72 hours post treatment, assay plates containing Vero cells were washed with Minimum Essential Medium (MEM, Gibco) twice and then stained with 1:10 mixture of AlamarBlue cell viability reagent (Invitrogen) and MEM. After two hours of incubation at 8% CO_2_ at 37°C, fluorescence intensity of each well was measured at 560nm absorbance and 590nm emission using Cytation 5 Plate reader (Biotek). Percentage viability was measured using this equation: *viability* % = 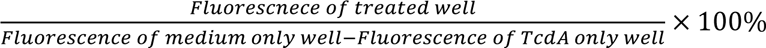. Experiments included three replicates whose standard deviation was calculated and reported as error bars (see Fig. 6b). Cytotoxicity assay was repeated more than three times.

## Figure generation

Graphics of cryo-EM density maps and molecular models were made using UCSF Chimera^43^ and PyMOL^49^. Figures were made using Adobe Illustrator and Microsoft Office. SEC, MALS, and cytotoxicity data were plotted using OriginLab.

## DATA AVAILABILITY

Cryo-EM density maps have been deposited in the EM databank (www.ebi.ac.uk/emdb) with the accession codes EMD-71326, EMD-71327, and EMD-71325 for hFcµ-tFcµ-chi3, hFcµ-tFcµ-chi4, and hFcµ respectively. The refined coordinate has been deposited in the Protein Data Bank (www.rcsb.org) with accession codes 9P6U, 9P6V, and 9P6T for hFcµ-tFcµ-chi3, hFcµ-tFcµ-chi4, and hFcµ respectively.

## AUTHOR CONTRIBUTIONS

The study was conceived by B.M.S and M.L.; experiments were conducted by M.L. and all authors contributed to data analysis and manuscript writing.

## ACKNOWLEGEMENTS

Cryogenic electron microscopy (cryo-EM) grids were screened at the Materials Research Laboratory Central Research Facilities, University of Illinois Champaign-Urbana with the assistance from Kristen M. Flatt. The cryo-EM data collection was performed at the Purdue Cryo-EM Facility with the assistance of Thomas Klose and Frank Vago. *C. difficile* experiments were conducted at facility provided by Michael Miller, using protocols developed with the help of Sonya Kumar Bharathkar and Minsoo Kim. This work was supported by NIH grant 1R01AI165570 and University of Illinois start-up funding to B.M.S, and the Lowell P. Hager Fellowship in Biochemistry to M.L.

## CONFLICT OF INTERESTS

B.M.S and M.L are listed as inventors on a provisional patent application that includes the design, production, and use of chimeric antibodies, some of which include teleost Ig heavy chain motifs.

## SUPPLEMENTARY FIGURES AND TABLES

**Supplementary Fig. 1.**
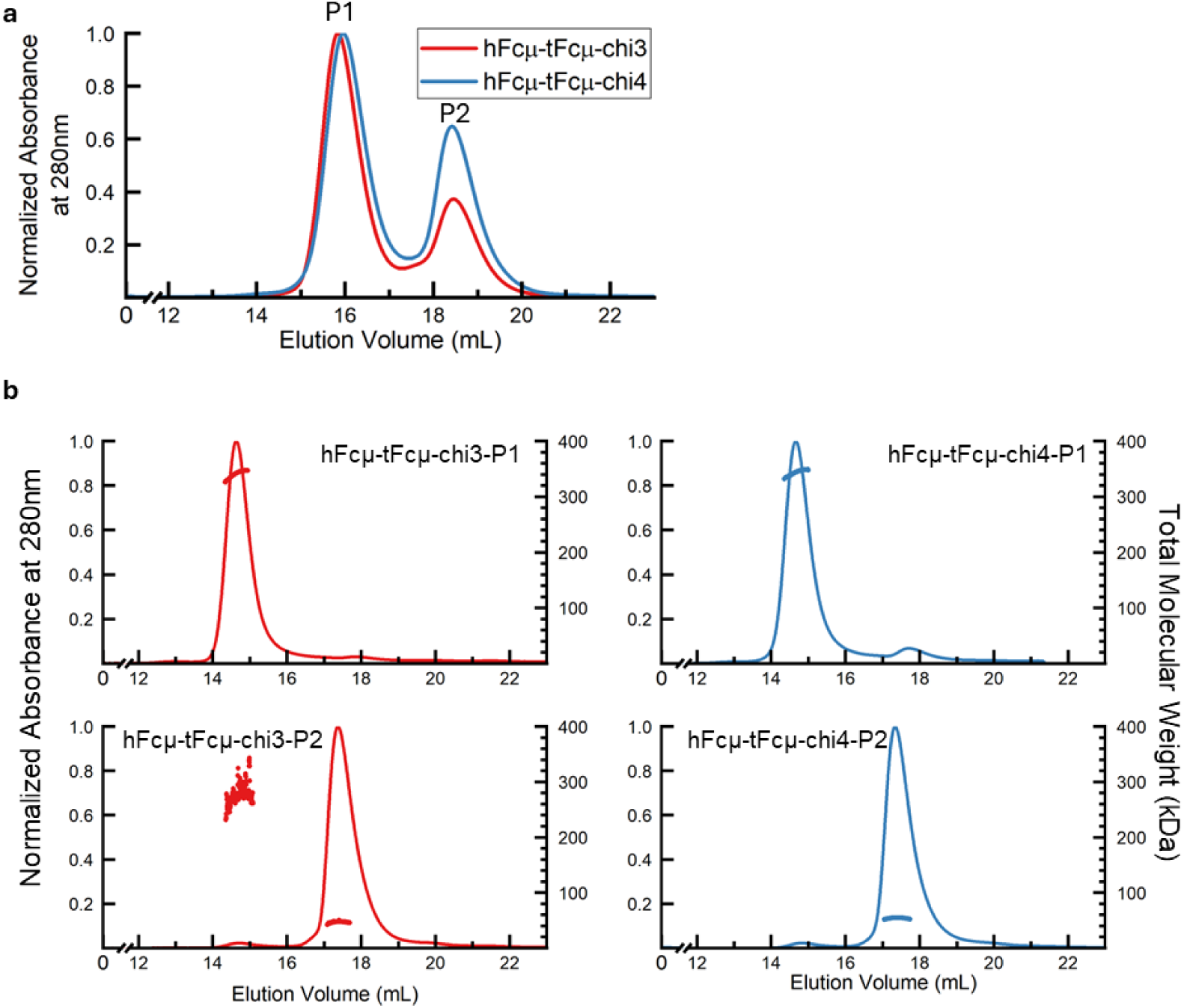
Purification and characterization of hFcμ-tFcμ-chi3 and hFcμ-tFcμ-chi4. **a** Superose 6 size exclusion column chromatograms of recombinantly expressed hFcμ-tFcμ-chi3 and hFcμ-tFcμ-chi4 post affinity chromatography purification. Each peak (P1 and P2) was pooled and analyzed using size exclusion chromatography with in-line multi-angle light scattering (SEC-MALS) **b** SEC-MALS plots for the indicated recombinant hFcμ-tFcμ-chi3 and hFcμ-tFcμ-chi4 complexes run on an Superdex 200 Increase column. For each panel, normalized absorbance at 280nm is shown on the left and total molecular weight (kDa) is shown on the right.

**Supplementary Fig. 2.**
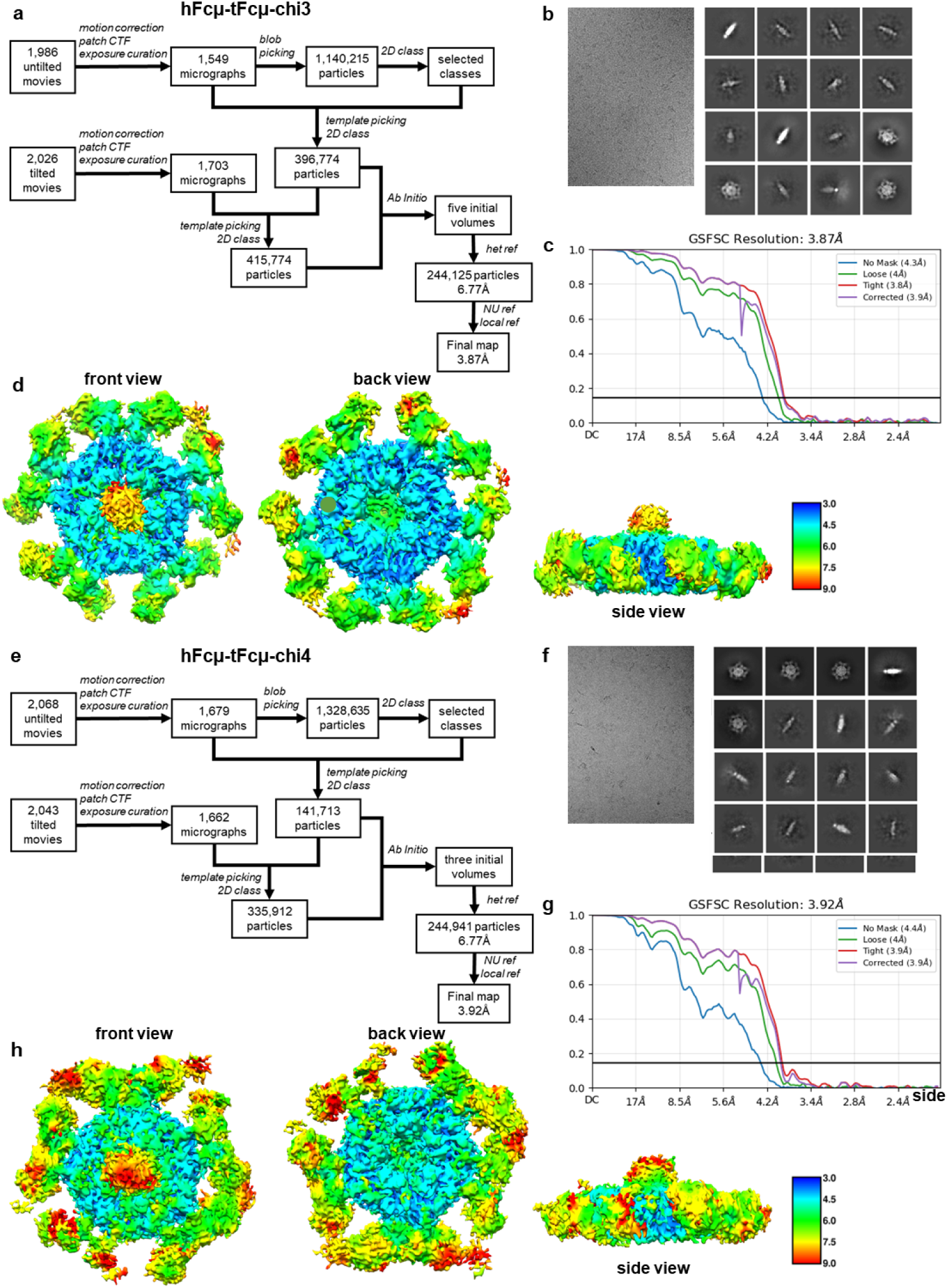
hFcµ-tFcµ-chi3 and hFcµ-tFcµ-chi4 cryo-EM data collection and processing pipeline. **a** Cryo-EM data processing pipeline for hFcµ-tFcµ-chi3 in CryoSPARC. **b** Representative micrograph and 2D class averages of hFcµ-tFcµ-chi3. **c** FSC curves for the final reconstruction of hFcµ-tFcµ-chi3. FSC = 0.143 is indicated by the blue horizontal line. **d** Local resolution map of the final hFcµ-tFcµ-chi3 reconstruction. Local resolution was calculated in CryoSparc and maps were rendered in UCSF Chimera. **e** Cryo-EM data processing pipeline for hFcµ-tFcµ-chi4 in CryoSPARC. **f** Representative micrograph and 2D class averages of hFcµ-tFcµ-chi4. **g** FSC curves for the final reconstruction of hFcµ-tFcµ-chi4. FSC = 0.143 is indicated by the blue horizontal line. **h** Local resolution map of the final hFcµ-tFcµ-chi4 reconstruction. Local resolution was calculated in CryoSparc and maps were rendered in UCSF Chimera. The unit of the scale bar is in Angstroms (Å)

**Supplementary Fig. 3.**
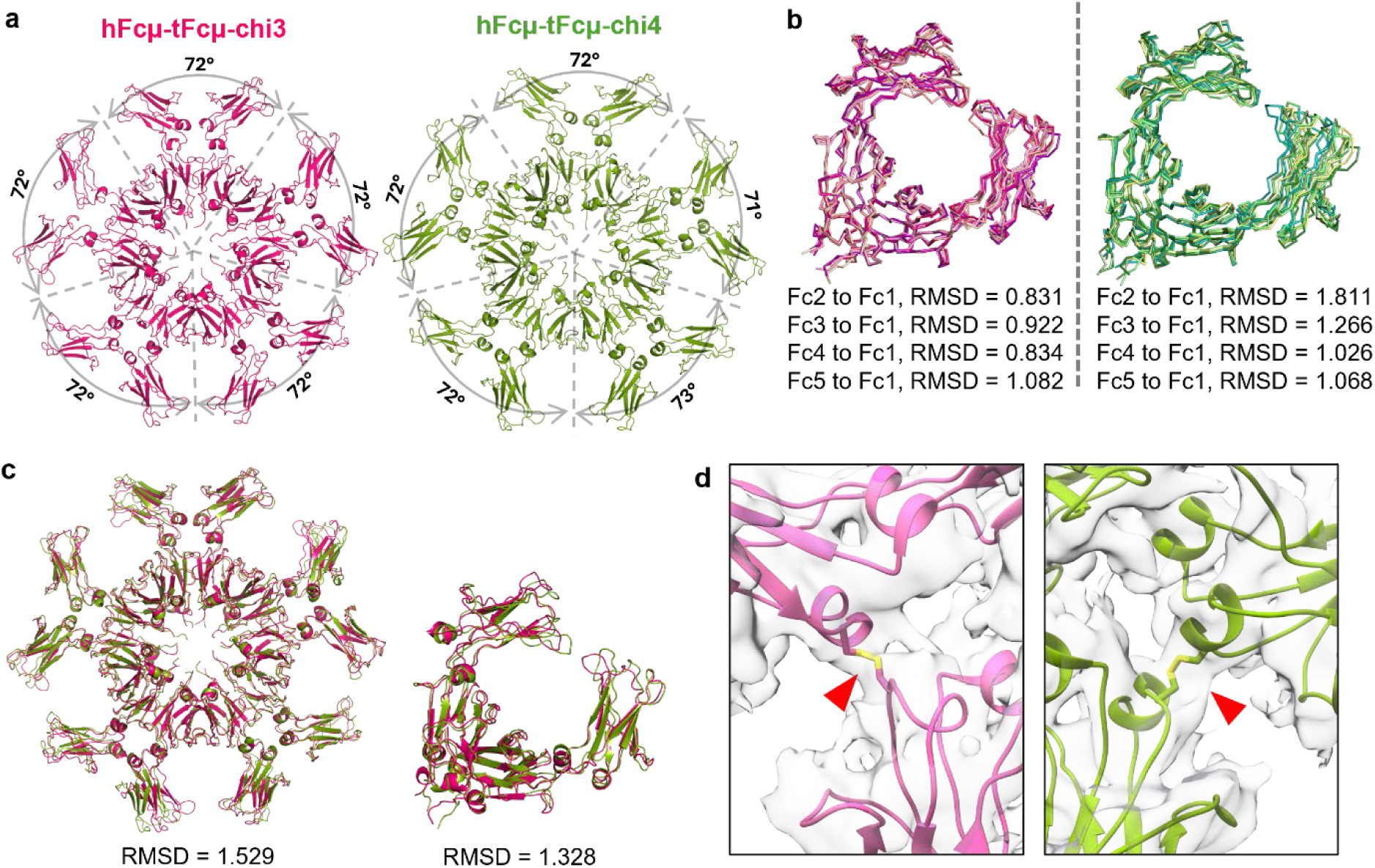
Structural alignments between hFcμ-tFcμ-chi3 and hFcμ-tFcμ-chi4. **a** Angles between adjacent Fcµ monomers for the hFcμ-tFcμ-chi3 pentamer (*left, pink*) and hFcμ-tFcμ-chi4 pentamer (*right, green*). **b** Alignment of all hFcμ-tFcμ-chi3 Fcµ monomers (*left*) and alignment all hFcμ-tFcμ-chi4 Fcµ monomers (*right*). **c** *Left*, alignment between hFcμ-tFcμ-chi3 pentamer and hFcμ-tFcμ-chi4 pentamer. *Right*, alignment between hFcμ-tFcμ-chi3 Fcµ1 and hFcμ-tFcμ-chi4 Fcµ1. **d** Cryo-EM density map (gray) and structure (cartoon representation) showing the region around interchain disulfides between Cys414 residues (sticks) on neighboring Fcs. Red triangles point to the density of disulfides.

**Supplementary Fig. 4.**
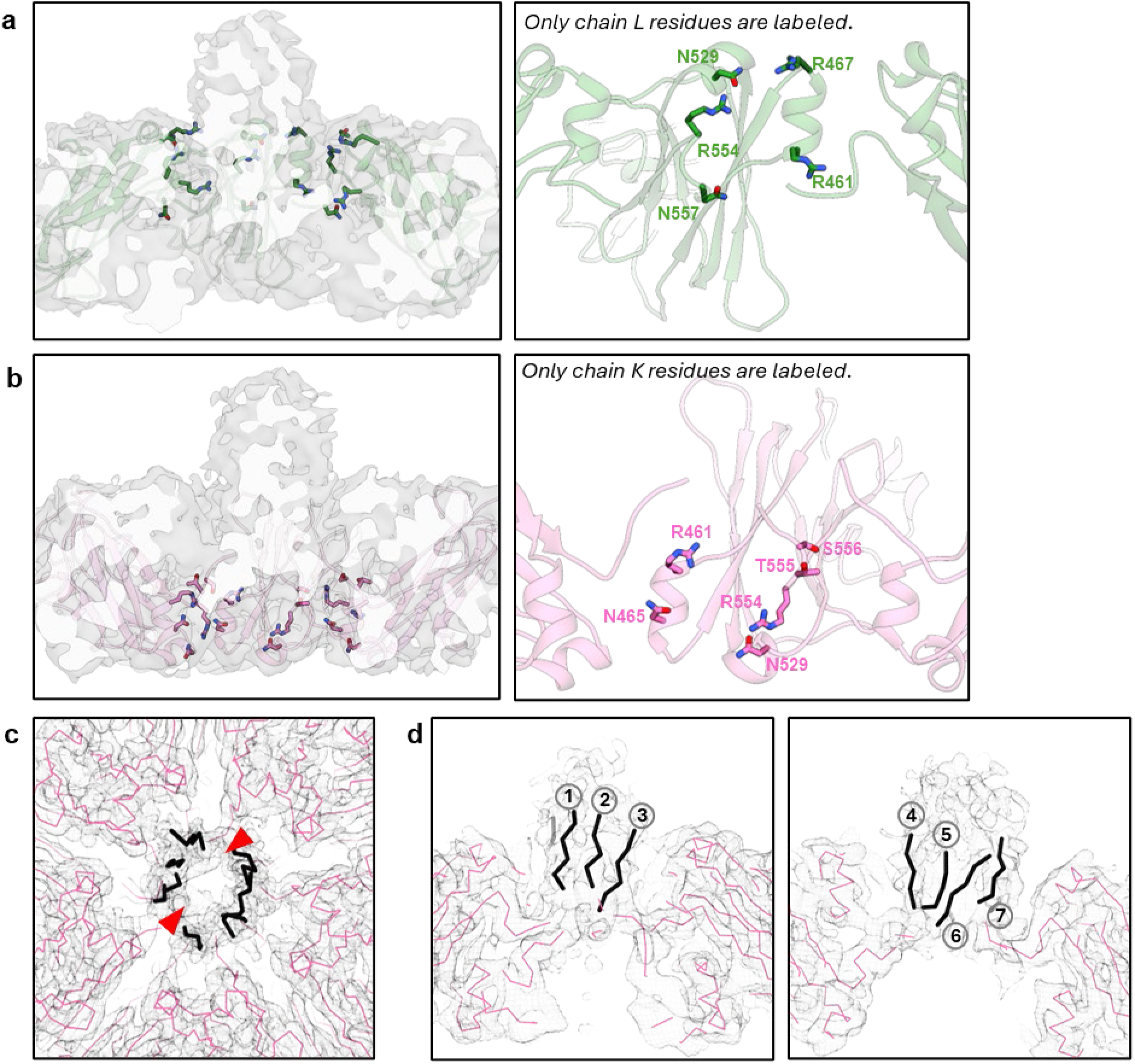
Molecular details of cFcµ CHμ4-pre-Tp interface, and Tp assembly model. **a** *Left*, hFcμ-tFcμ-chi3 density map with cartoon representation including the pre-Tp loops of chains B, H, L with residues Arg461, Arg467, Asn529, Arg554, Asn557 shown as sticks. *Right*, close-up view of chain L with the same residues shown in sticks as the left panel. **b** *Left*, hFcμ-tFcμ-chi3 density map with cartoon representation including the pre-Tp loops of chains A, G, K with residues Arg461, Arg467, Asn529, Arg554, Thr555, Ser556 shown as sticks. *Right*, Close-up view of chain K with the same residues shown in sticks as the left panel. **c** Top-down view of the solvent accessible tunnel at the center of the Tp assembly (indicated by red triangles). **d** hFcμ-tFcμ-chi3 density map and ribbon representation (pink) shown in two orientations with seven poly-alanine β-strands (black) fit to the Tp protrusion density. These strands were not included in the refined molecular model deposited in the PDB.

**Supplementary Fig. 5.**
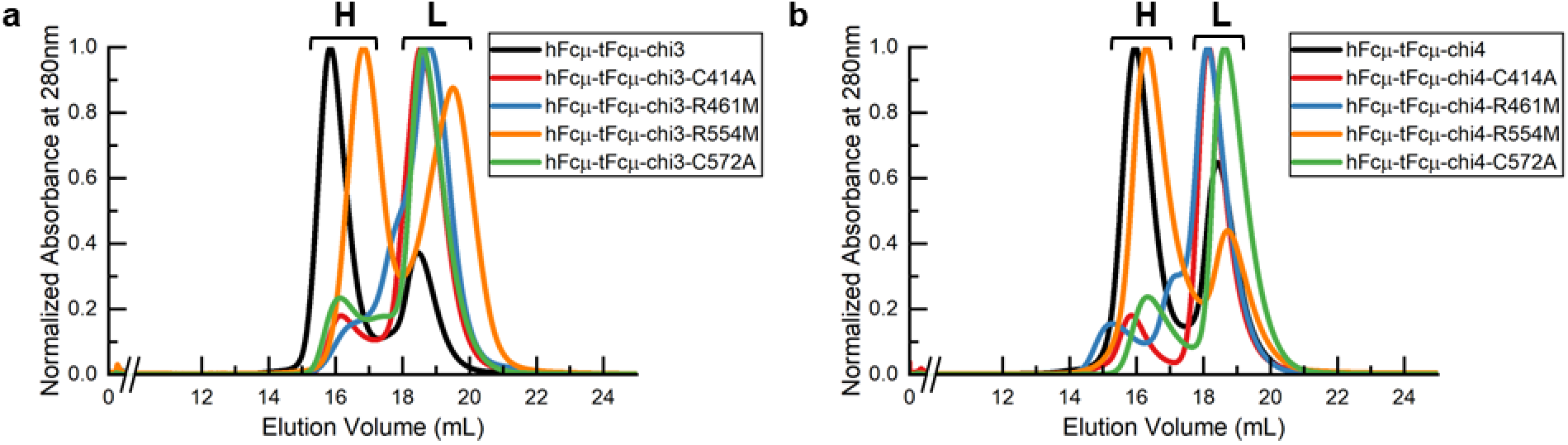
SEC chromatographs of hFcμ-tFcμ-chi3, hFcμ-tFcμ-chi4, and mutant variants. **a** SEC chromatographs of hFcμ-tFcμ-chi3 and the indicated mutant variants and **b** SEC chromatographs of hFcμ-tFcμ-chi4 and the indicated single mutant variants. “H” stands for high molecular weight peak and “L” stands for low molecular weight peak. Results are consistent with SEC analysis that has been repeated at least twice for each sample (not shown).

**Supplementary Fig. 6.**
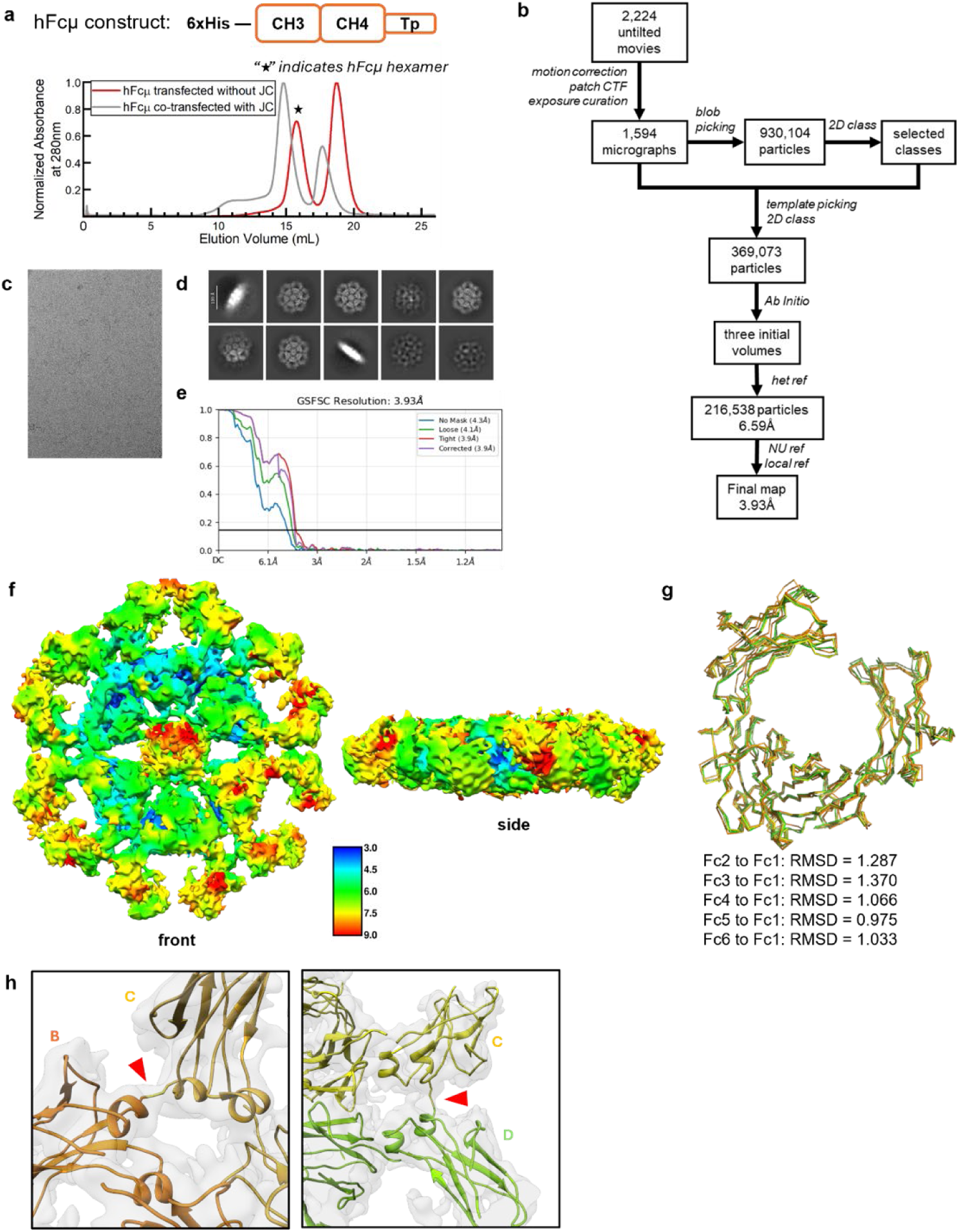
hFcμ hexamer purification, cryo-EM data collection and processing pipeline, and structural analysis. **a** SEC chromatographs of purified hFcμ resulting from transfection without JC and purified Fcμ-J resulting from transfection with JC. The JC-free hFcμ hexamer (indicated by star) was used for subsequence cryo-EM data collection. **b** Schematic summary of hFcμ hexamer cryo-EM data processing pipeline in CryoSPARC. **c** Representative micrograph after motion correction and selection. **d** Representative 2D class averages. **e** FSC curves for the final reconstruction with reported resolution at FSC = 0.143 shown by the blue horizontal line. **f** Local resolution maps of the final reconstruction calculated in CryoSPARC and rendered in UCSF Chimera; the unit of scale bar is in Angstroms (Å). **g** Alignment between hFcμ monomers within the structure. **h** Cryo-EM density map (gray) and structure (cartoon) showing interchain disulfides formed between Cys414 residues (sticks) on neighboring Fcμs. Red triangles point to the density of disulfides.

**Supplementary Fig. 7.**
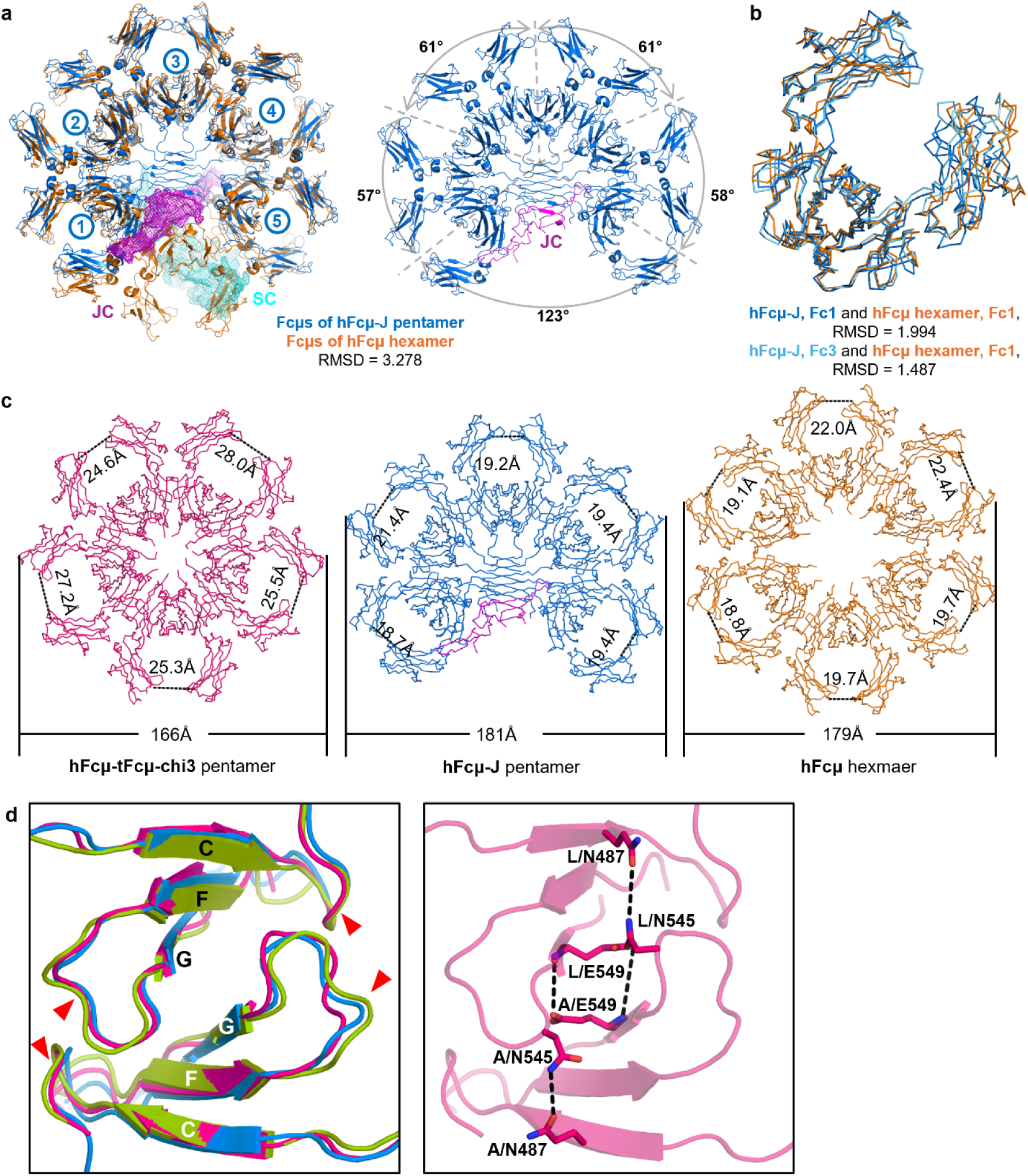
Structural comparison between cFcμs, hFcμ-J pentamer, and hFcμ hexamer. **a** Structural alignment between pentameric SIgM structure (hFcμ-J and SC; PDB ID: <u>6KXS</u>) and hFcμ hexamer. All chains were included in the alignment. **b** Structural alignment between Fcμ3 of Fcμ-J pentamer (blue) and Fcμ1 of hFcμ hexamer (orange). **c** All three structures are shown as ribbons with the CHμ3-CHμ3 distance within each Fcμ monomer and the overall diameter of each structure indicated. **d** *Left*, Fc-Fc interface in hFcμ-tFcμ-chi3 (pink), hFcμ-tFcμ-chi4 (green), and hIgM-J pentamer (*blue*), with β-strands C, F, G labeled. Red triangles point to CD loops and FG loops. *Right*, Fcμ-Fcμ interface of hFcμ-tFcμ-chi3 with residues contributing to polar contacts shown as sticks.

**Supplementary Fig. 8.**
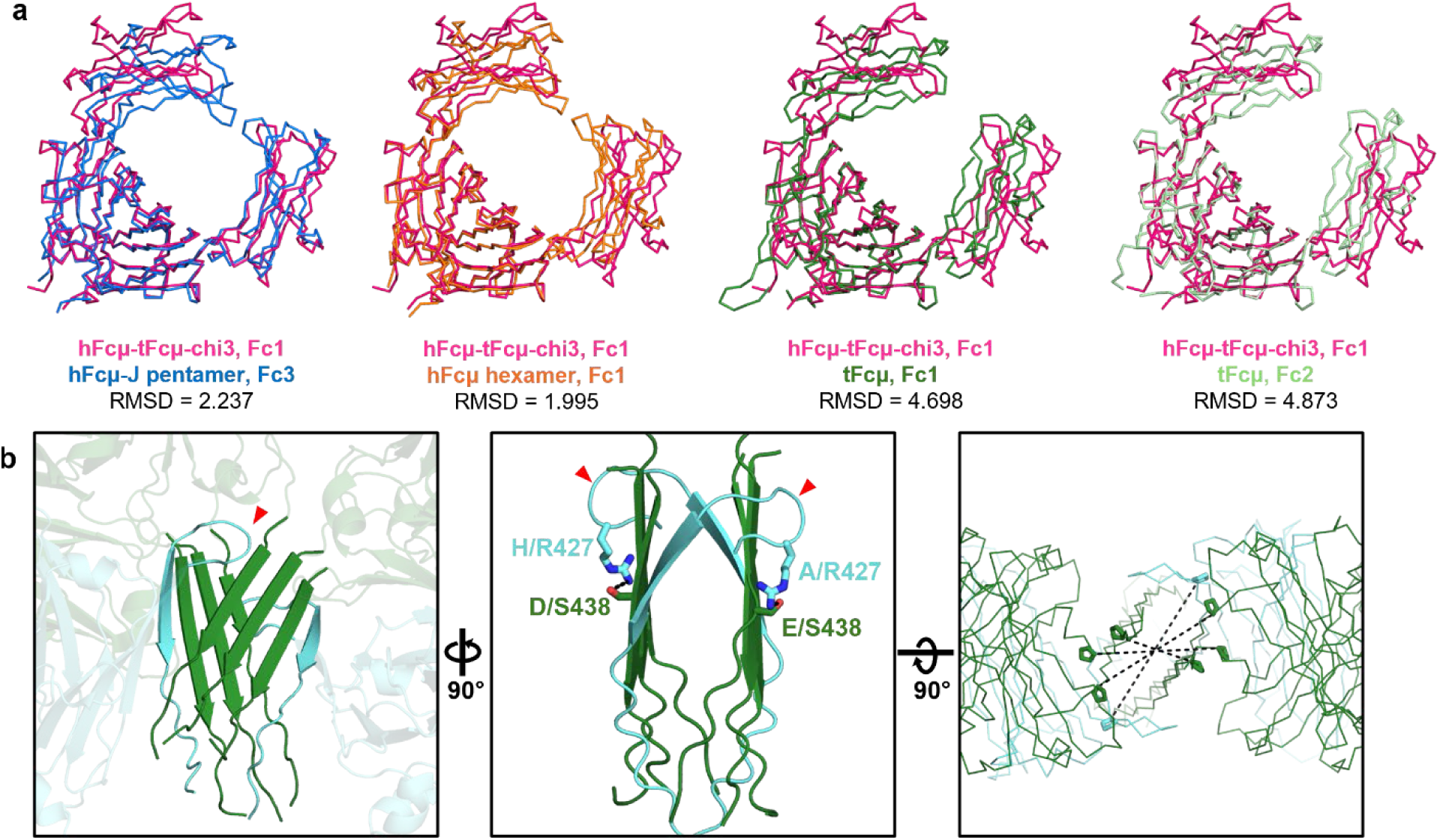
Structural alignment between individual Fcμs and the structure of the tFcμ tetramer Tp assembly. **a** Pairwise alignments of the indicated Fcμ monomers colored according to the key below each Fcμ pair. **b** Close-up view of the tFcμ tetramer Tp assembly shown from three perspectives (PDB ID: <u>8GHZ</u>). Chains A and H are colored cyan and other chains are colored green. Red triangles point to pre-Tp loops formed by chains A and H. *Left*, front view of Tp assembly. *Middle*, side view of Tp assembly. Residues contributing to interactions between pre-Tp loops and Tp residues are shown in sticks. *Right*, top-down view of Tp assembly with Pro432 shown as sticks.

**Supplementary Fig. 9.**
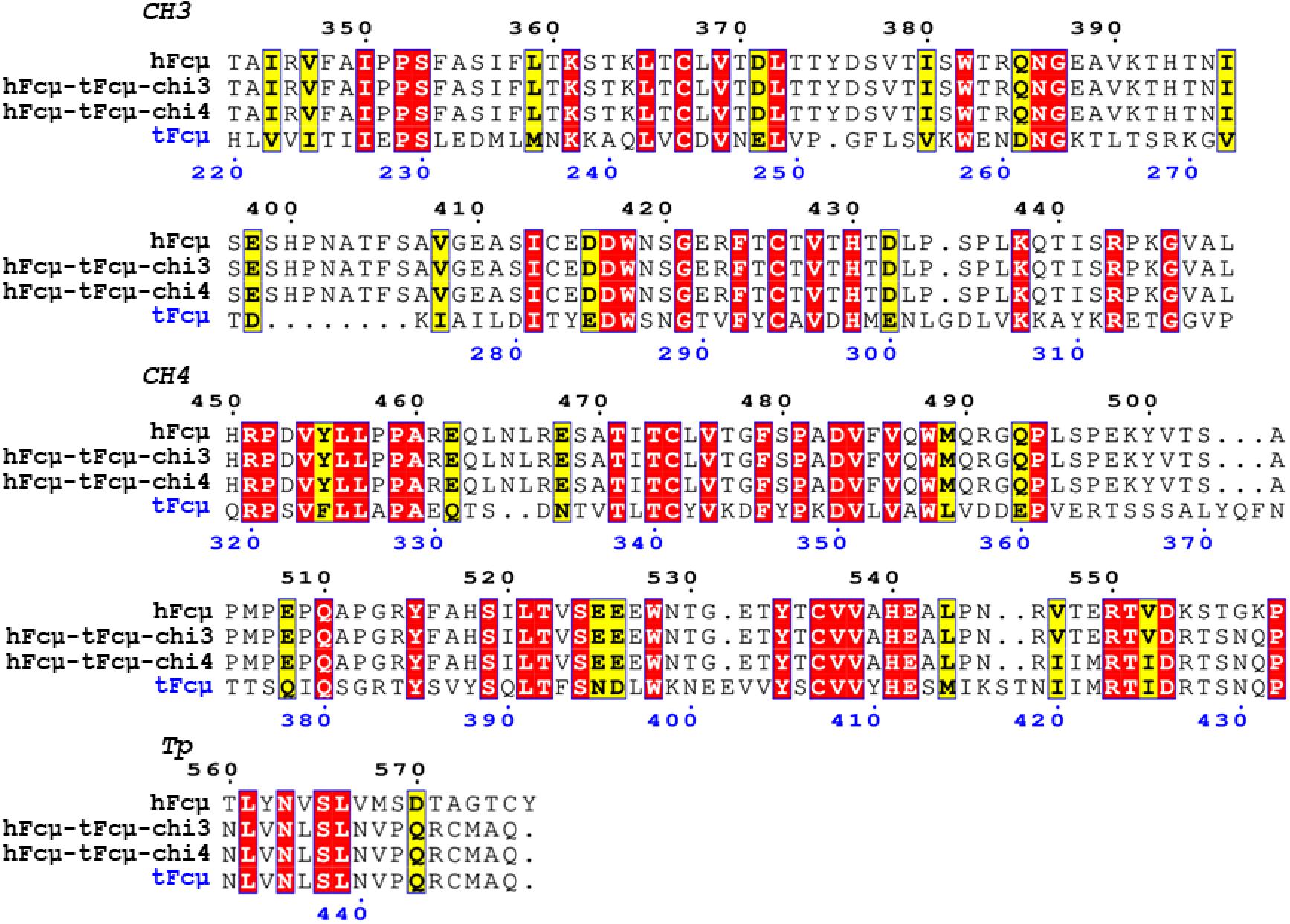
Sequence alignment between hFcμ, tFcμ, and cFcμ sequences. Residue numbering of hFcμ and cFcμs is indicated above the sequences in black text and residue numbering of tFcμ is indicated below sequences in blue text. Residues are colored based on similarity score; red boxes indicate conserved residues and yellow boxes indicate similar residues.

**Supplementary Table 1.**
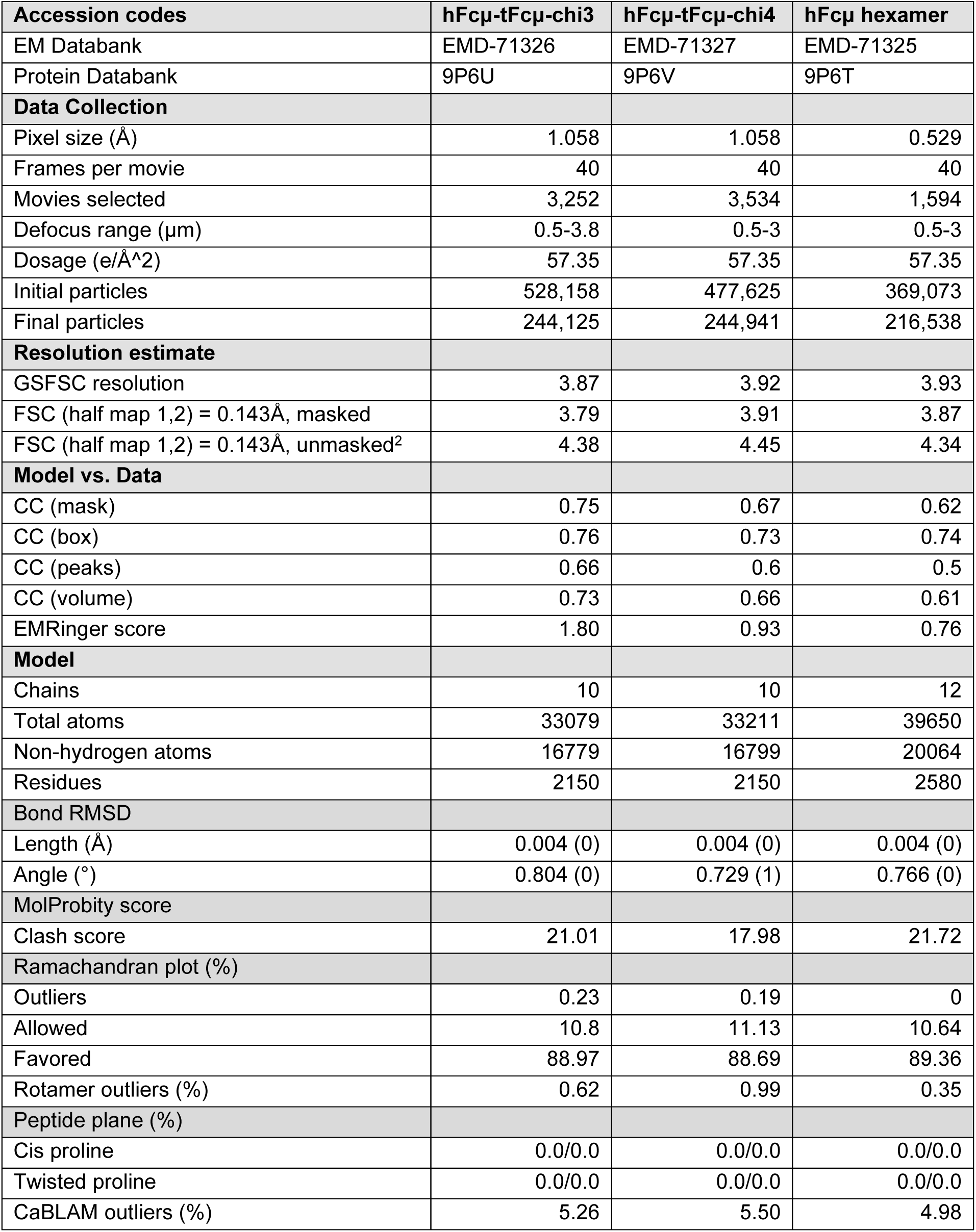
Cryo-EM data collection and refinement statistics associated with hFcμ-tFcμ-chi3, hFcμ-tFcμ-chi4, and hFcμ. Map resolution was estimated using CryoSPARC and PHENIX. Models were refined and validated using Coot and PHENIX.

## REFERENCES

1. Keyt, B. A., Baliga, R., Sinclair, A. M., Carroll, S. F. & Peterson, M. S. Structure, Function, and Therapeutic Use of IgM Antibodies. Antibodies 9, 53 (2020).

2. Flajnik, M. F. A cold-blooded view of adaptive immunity. Nat Rev Immunol 18, 438–453 (2018).

3. Eve, O., Matz, H. & Dooley, H. Proof of long-term immunological memory in cartilaginous fishes. Developmental & Comparative Immunology 108, 103674 (2020).

4. Pettinello, R. & Dooley, H. The Immunoglobulins of Cold-Blooded Vertebrates. Biomolecules 4, 1045–1069 (2014).

5. Brewer, J. W., Randall, T. D., Parkhouse, R. M. E. & Corley, R. B. IgM hexamers? Immunology Today 15, 165–168 (1994).

6. Eskeland, T. & Christensen, T. B. IgM Molecules with and without J Chain in Serum and after Purification, Studied by Ultra-centrifugation, Electrophoresis, and Electron Microscopy. Scand J Immunol 4, 217–228 (1975).

7. Wiersma, E. J. & Shulman, M. J. Assembly of IgM. Role of disulfide bonding and noncovalent interactions. J Immunol 154, 5265–5272 (1995).

8. Boes, M. Role of natural and immune IgM antibodies in immune responses. Molecular Immunology 37, 1141–1149 (2000).

9. Oskam, N. et al. At Critically Low Antigen Densities, IgM Hexamers Outcompete Both IgM Pentamers and IgG1 for Human Complement Deposition and Complement-Dependent Cytotoxicity. The Journal of Immunology 209, 16–25 (2022).

10. Kaetzel, C. S. The polymeric immunoglobulin receptor: bridging innate and adaptive immune responses at mucosal surfaces. Immunol Rev 206, 83–99 (2005).

11. Kong, X. et al. Comparison of polymeric immunoglobulin receptor between fish and mammals. Veterinary Immunology and Immunopathology 202, 63–69 (2018).

12. Chen, K., Magri, G., Grasset, E. K. & Cerutti, A. Rethinking mucosal antibody responses: IgM, IgG and IgD join IgA. Nat Rev Immunol 20, 427–441 (2020).

13. Kaetzel, C. S. Coevolution of Mucosal Immunoglobulins and the Polymeric Immunoglobulin Receptor: Evidence That the Commensal Microbiota Provided the Driving Force. ISRN Immunology 2014, 1–20 (2014).

14. Rochereau, N. et al. Essential role of TOSO/FAIM3 in intestinal IgM reverse transcytosis. Cell Reports 37, 110006 (2021).

15. Li, Y. et al. Structural insights into immunoglobulin M. Science 367, 1014–1017 (2020).

16. Kumar, N., Arthur, C. P., Ciferri, C. & Matsumoto, M. L. Structure of the human secretory immunoglobulin M core. Structure 29, 564–571.e3 (2021).

17. Chen, Q., Menon, R., Calder, L. J., Tolar, P. & Rosenthal, P. B. Cryomicroscopy reveals the structural basis for a flexible hinge motion in the immunoglobulin M pentamer. Nat Commun 13, 6314 (2022).

18. Müller, R. et al. High-resolution structures of the IgM Fc domains reveal principles of its hexamer formation. Proc. Natl. Acad. Sci. U.S.A. 110, 10183–10188 (2013).

19. Hiramoto, E. et al. The IgM pentamer is an asymmetric pentagon with an open groove that binds the AIM protein. Sci. Adv. 4, eaau1199 (2018).

20. Lyu, M., Malyutin, A. G. & Stadtmueller, B. M. The structure of the teleost Immunoglobulin M core provides insights on polymeric antibody evolution, assembly, and function. Nat Commun 14, 7583 (2023).

21. Pasalic, D. et al. A peptide extension dictates IgM assembly. Proc Natl Acad Sci USA 114, E8575–E8584 (2017).

22. Kaattari, S. et al. Varied redox forms of teleost IgM: an alternative to isotypic diversity? Immunol Rev 166, 133–142 (1998).

23. Cattaneo, A. & Neuberger, M. S. Polymeric immunoglobulin M is secreted by transfectants of non-lymphoid cells in the absence of immunoglobulin J chain. The EMBO Journal 6, 2753–2758 (1987).

24. Randall, T. D., Brewer, J. W. & Corley, R. B. Direct evidence that J chain regulates the polymeric structure of IgM in antibody-secreting B cells. Journal of Biological Chemistry 267, 18002–18007 (1992).

25. Niles, M. J., Matsuuchi, L. & Koshland, M. E. Polymer IgM assembly and secretion in lymphoid and nonlymphoid cell lines: evidence that J chain is required for pentamer IgM synthesis. Proc. Natl. Acad. Sci. U.S.A. 92, 2884–2888 (1995).

26. Chen, Q., Menon, R. P., Masino, L., Tolar, P. & Rosenthal, P. B. Structural basis for Fc receptor recognition of immunoglobulin M. Nat Struct Mol Biol 30, 1033–1039 (2023).

27. Li, Y. et al. Immunoglobulin M perception by FcμR. Nature 615, 907–912 (2023).

28. Hussack, G. et al. Neutralization of Clostridium difficile Toxin A with Single-domain Antibodies Targeting the Cell Receptor Binding Domain. Journal of Biological Chemistry 286, 8961–8976 (2011).

29. Chandrasekaran, R. & Lacy, D. B. The role of toxins in Clostridium difficile infection. FEMS Microbiology Reviews 41, 723–750 (2017).

30. Markantonis, J. E., Fallon, J. T., Madan, R. & Alam, M. Z. Clostridioides difficile Infection: Diagnosis and Treatment Challenges. Pathogens 13, 118 (2024).

31. Bharathkar, S. K., Miller, M. J. & Stadtmueller, B. M. Engineered Secretory Immunoglobulin A provides insights on antibody-based effector mechanisms targeting Clostridiodes difficile. Preprint at 10.1101/2023.11.08.566291 (2023).

32. Anosova, N. G. et al. A Combination of Three Fully Human Toxin A- and Toxin B-Specific Monoclonal Antibodies Protects against Challenge with Highly Virulent Epidemic Strains of Clostridium difficile in the Hamster Model. Clin. Vaccine Immunol. 22, 711–725 (2015).

33. Kumar, N., Arthur, C. P., Ciferri, C. & Matsumoto, M. L. Structure of the secretory immunoglobulin A core. Science 367, 1008–1014 (2020).

34. Kumar Bharathkar, S., et al. The structures of secretory and dimeric immunoglobulin A. eLife 9, e56098 (2020).

35. Ye, J., Bromage, E. S. & Kaattari, S. L. The Strength of B Cell Interaction with Antigen Determines the Degree of IgM Polymerization. The Journal of Immunology 184, 844–850 (2010).

36. Giannone, C. et al. Biogenesis of secretory immunoglobulin M requires intermediate non-native disulfide bonds and engagement of the protein disulfide isomerase ERp44. The EMBO Journal 41, e108518 (2022).

37. Matsumoto, M. L. Molecular Mechanisms of Multimeric Assembly of IgM and IgA. Annu. Rev. Immunol. 40, 221–247 (2022).

38. Xiang, Y. et al. Adaptive multi-epitope targeting and avidity-enhanced nanobody platform for ultrapotent, durable antiviral therapy. Cell 187, 6966–6980.e23 (2024).

39. Murase, T. et al. Structural Basis for Antibody Recognition in the Receptor-binding Domains of Toxins A and B from Clostridium difficile. Journal of Biological Chemistry 289, 2331–2343 (2014).

40. Punjani, A., Rubinstein, J. L., Fleet, D. J. & Brubaker, M. A. cryoSPARC: algorithms for rapid unsupervised cryo-EM structure determination. Nat Methods 14, 290–296 (2017).

41. Punjani, A., Zhang, H. & Fleet, D. J. Non-uniform refinement: adaptive regularization improves single-particle cryo-EM reconstruction. Nat Methods 17, 1214–1221 (2020).

42. Waterhouse, A. et al. SWISS-MODEL: homology modelling of protein structures and complexes. Nucleic Acids Research 46, W296–W303 (2018).

43. Pettersen, E. F. et al. UCSF Chimera--A visualization system for exploratory research and analysis. J. Comput. Chem. 25, 1605–1612 (2004).

44. Emsley, P., Lohkamp, B., Scott, W. G. & Cowtan, K. Features and development of *Coot*. Acta Crystallogr D Biol Crystallogr 66, 486–501 (2010).

45. Liebschner, D. et al. Macromolecular structure determination using X-rays, neutrons and electrons: recent developments in *Phenix*. Acta Crystallogr D Struct Biol 75, 861–877 (2019).

46. Barad, B. A. et al. EMRinger: side chain–directed model and map validation for 3D cryo-electron microscopy. Nat Methods 12, 943–946 (2015).

47. Sievers, F. et al. Fast, scalable generation of high-quality protein multiple sequence alignments using Clustal Omega. Mol Syst Biol 7, 539 (2011).

48. Robert, X. & Gouet, P. Deciphering key features in protein structures with the new ENDscript server. Nucleic Acids Research 42, W320–W324 (2014).

49. Schrödinger, LLC. The PyMOL Molecular Graphics System, Version 3.0. (2015).

